# Evaluating effects of elevated [CO_2_] and accelerated NPQ relaxation on yield, physiology and transcription in soybean

**DOI:** 10.1101/2025.11.19.689310

**Authors:** Dilkaran Singh, Amanda P. De Souza, Lynn Doran, Jeffrey Hansen, Steven J. Burgess

**Affiliations:** Carl R Woese Institute for Genomic Biology, University of Illinois Urbana-Champaign; Department of Plant Biology, University of Illinois Urbana-Champaign

**Author notes:** these authors contributed equally to the manuscript (Dilkaran Singh, Amanda P. De Souza).

## Abstract

Improving photosynthetic efficiency is proposed as a means of enhancing crop productivity under rising atmospheric CO_2_ concentrations ([CO_2_]). To assess whether additive gains can be realized from simultaneous improvements in light use-efficiency and carbon assimilation, we investigated the physiological and molecular responses to elevated CO_2_ (e[CO_2_]) of a transgenic soybean line (VPZ-34A) expressing the genes violaxanthin de-epoxidase (*VDE*), Photosystem II subunit S (*PsbS*) and zeaxanthin epoxidase (*ZEP*) from *Arabidopsis thaliana* (arabidopsis) (*AtVPZ*). Plants were grown at a free-air CO_2_ enrichment facility (SoyFACE) in 2021, with physiological and transcriptomic measurements collected during vegetative (V4-5) and reproductive stages (R5-6), then compared to final biomass and seed production parameters. VPZ-34A showed increased maximum quantum photosynthetic efficiency of photosynthesis (φPSII_max_) under fluctuating light during vegetative stages, but no increase in maximum efficiency of carbon assimilation (φCO_2, max_), and no significant increase in seed or biomass related traits under ambient or e[CO_2_]. Transcriptome analysis showed the VPZ construct impacted a small number of genes, among them downregulation of a mitochondrial thioredoxin (*Trx o1*) and up-regulation of two *AP2* transcription factors in VPZ lines may suggest alterations in redox balance and seed development. In conclusion, the improvements in φPSII_max_ and NPQ in VPZ-34A were unaffected by e[CO_2_] but didn’t translate to stimulation of yield. Our data highlight the complexity of VPZ manipulation on plant performance and provide avenues for future research to optimize this strategy to improve crop yields.

## INTRODUCTION

Approximately 733 million people faced hunger in 2023, and this number has risen since 2019 (FAO 2024). Changes in climate, natural disasters, and armed conflicts further threaten global food security (Oladipo and Oyinloye 2022; FAO 2024; Iriti and Vitalini 2025). Soybean is the most widely cultivated legume with more than 371 million tons produced worldwide in 2023, making it a key crop for food security perspective (FAO 2025). Threat of the aforementioned challenges have driven the development of numerous biotechnological approaches to improve crop yield, including soybean (Anderson et al. 2016; Munaweera et al. 2022; Tyczewska et al. 2023). One of the strategies that has promising results is the improvement of photosynthesis (Simkin et al. 2015; Kromdijk et al. 2016; South et al. 2019; López-Calcagno et al. 2020; De Souza et al. 2022). However, converting improvements at leaf level photosynthesis to crop yield is not straightforward. For example, elevated [CO_2_] (e[CO_2_]) is known to enhance light saturated photosynthesis in C_3_ plants by increasing Rubisco carboxylation efficiency and reducing photorespiration through competitive inhibition of oxygenation (Drake et al. 1997; Ainsworth and Long 2005; Bernacchi et al. 2005). However, in soybean, the intensity of the photosynthetic response to e[CO_2_] was shown to be strongly associated with the yield potential of the cultivar evaluated (Ainsworth and Long 2021), highlighting the importance of sink strength. Further, a cross-scale mathematical model shows that the yield benefit from increasing photosynthetic rates can vary depending on the environmental conditions (Wu et al. 2023). These findings indicate a greater understanding of how manipulations of photosynthesis interact with whole plant physiology are necessary for the success of engineering approaches.

Several studies have looked to improve the efficiency of light use under fluctuating light (Kromdijk et al. 2016; Garcia-Molina and Leister 2020; Zhu 2021; De Souza et al. 2022; Lehretz et al. 2022; Milburn et al. 2025). When the supply of photons exceeds demand, such as during high light conditions or drought, build-up of reduced electron transfer intermediates in the photosynthetic electron transport chain can initiate the production of reactive oxygen species, leading to photodamage and a reduction in photosynthetic activity (Anderson et al. 1998; Mellis 1999; Rutherford et al. 2012). A variety of mechanisms, collectively referred to as non-photochemical quenching of chlorophyll fluorescence excited states (NPQ), protect leaves against photodamage by dissipating light energy as heat (Niyogi and Truong 2013; Ruban 2016; Zuo 2025). When light is in excess, protective NPQ is beneficial, and it has been shown to be essential for plant fitness under fluctuating light conditions (Külheim et al. 2002). However, when leaves transition from full sun to shade, energy dissipation by NPQ can be slow to adjust, and under such transitory conditions, NPQ can divert energy away from photosynthesis (Zhu et al. 2004; Murchie and Niyogi 2011). This is important, as slow relaxation of NPQ in soybean is predicted to cause an 11% reduction in maximum theoretical daily carbon assimilation (Wang et al. 2020).

Light induces proton pumping across the thylakoid membrane leading to lumen acidification and build-up of a proton motive force (*pmf*), which is used to drive ATP synthesis. However, NPQ is also triggered by lumen acidification (Wraight and Crofts 1970; Briantais et al. 1979), and acts as a feedback mechanism, modulating light harvesting capacity according to energetic demand. The precise mechanisms of NPQ are debated (for review see (Ruban 2016)), but, fast energy-dependent quenching “qE” has been shown to be related to acidification of the lumen, triggering a conformational change of PSII-associated antennae via protonation of Photosystem II (PSII) subunit S (PsbS), leading to quenching (Li et al. 2000; Johnson et al. 2011; Wilson et al. 2024). In addition, lumen acidification activates violaxanthin de-epoxidase (VDE), which catalyzes the conversion of violaxanthin (Vx) to zeaxanthin (Zx) via antheraxanthin (Yamamoto et al. 1962; Hager 1969; Yamamoto and Kamite 1972), with accumulation of Zx involved in modulation of quenching (Demmig Adams et al. 1996; Niyogi et al. 1998). An intermediate relaxing component of NPQ, termed “qZ” is associated with the catabolism of zeaxanthin, and typically operates on the time scale of 10-15 mins (Niyogi et al. 1998; Nilkens et al. 2010). This conversion of Zx back to Vx, is catalyzed by the stromal localized enzyme zeaxanthin epoxidase (ZEP) (Sapozhnikov 1957; Siefermann and Yamamoto 1975; Marin et al. 1996) and is involved in down-regulating NPQ.

Following model predictions, Kromdijk et al. (2016) overexpressed homologs of *VDE*, *PsbS* and *ZEP* genes from arabidopsis (*AtVPZ*) in tobacco, to speed up relaxation of NPQ and reduce the loss of potential energy during shadeflecks. This was found to lead to an increase in light use efficiency under fluctuating light, which translated to greater biomass production in small scale field experiments (Kromdijk et al. 2016). A follow-up study in soybean applied the same strategy and observed accelerated response of the fast (τ_1_; qE) and intermediate (τ_2_; qZ) time constants for NPQ relaxation upon transition from sun to shade by 37% and 67%, respectively (De Souza et al. 2022). This resulted in an improvement in photosynthetic efficiency under fluctuating light and an average increase of 24.5% in seed production compared to untransformed soybean in one out of two years of experiments (De Souza et al. 2022). Similar efforts in other species have resulted in reduced biomass and/or yield, including arabidopsis (Garcia-Molina and Leister 2020), potato (Lehretz et al. 2022), and creeping bentgrass (Zhu 2021), whereas in setaria improvement in yield under specific conditions was observed (Milburn et al. 2025). Therefore, the particular conditions under which this modification is likely to be beneficial are not clearly understood, necessitating further investigation.

A variety of modeling approaches have assessed the potential impact of combining improvements in CO_2_ fixation and light reactions. Typically, these models predict an additive effect, although the precise scale of improvement and influence of environment vary (Wu et al. 2023). Improved response from additive effects also find some support from experiments in tobacco involving overexpression of SBPase and an algal cytochrome b_6_f protein (López-Calcagno et al. 2020). Given the promising findings of De Souza et al. (2022), we sought to test whether stacking increases in light use efficiency with increased CO_2_ fixation would lead to additive gains. Therefore, we characterized the physiological and transcriptional responses of a wild-type and a transgenic VPZ soybean line, under field conditions, comparing growth under elevated [CO_2_] levels as a proxy for increased carbon assimilation.

## MATERIALS AND METHODS

### Field site, plant material, and experimental design

This study was conducted during the summer of 2021 at the Soybean Free-Air CO_2_ Enrichment (SoyFACE) facility in Champaign, IL, USA (https://soyface.illinois.edu/) as described in (Aspray et al. 2023). *Glycine max* cv. Maverick; ND-18-34A, (hereafter named VPZ-34A) overexpressing three genes from *A. thaliana*: violaxanthin de-epoxidase (*AtVDE),* photosystem II subunit S (*AtPsbS*), and zeaxanthin epoxidase (*AtZEP*) was used in this study. This transgenic line was selected due to its higher seed production in previous experiments (De Souza et al. 2022). T5 seeds from the transgenic line were obtained as previously described (De Souza et al. 2022) and planted paired with a control of non-transgenic seeds (WT) from the same cultivar.

The experiment had four experimental blocks. Each block consisted of one plot that was fumigated to elevate the [CO_2_] to 600 µmol mol^-1^ paired with another plot that served as a control with the current a[CO_2_] concentration of 410 µmol mol^-1^. Each plot (7 m^2^) was split in two sub-plots to accommodate four rows of 1.2 m for each one of the lines. Each row had 34 seeds, totaling 136 plants per sub-plot. Row spacing was 0.75 m, and seed spacing 3.8 cm. Seed planting depth was 3.5 cm. The experiment had a 10 m bare border to comply with APHIS/USDA regulatory permits. All the measurements and tissue sampling were done in the middle two rows of each sub-plot.

A few days before the planting, the field was cultivated and tilled. On June 8^th^, 2021, T5 seeds from VPZ-34A and WT were hand-planted. Prior to planting, seeds were treated with CruiserMaxx Vibrance^®^ to protect against damage from early-season insects and diseases. Diatomaceous earth (PF Harris®, Cartersville, GA, USA) was added 8 days after sowing (DAS) to prevent cutworms. For pest control, the broad-spectrum insecticide Mustang^®^ Maxx (FMC Corporation., Philadelphia, PA, USA) was applied during the V4-V5 stage following the manufacturer’s instructions. To allow homogeneous germination, irrigation was supplied manually until the emergence of seedlings. After that stage, the experiment was rainfed.

### Measurement of NPQ

The speed of NPQ relaxation during the transition from high light to low light was assessed using a chlorophyll fluorescence imager (CF Imager, Technologica Ltd., UK) following the protocol described by (Gotarkar et al. 2022). Leaf disks were sampled at 32 DAS during V3-V4 stage. Time constants related to qE (τ_1_), qZ (τ_2_), and NPQ at steady-state high light (NPQ_max)_ were calculated fitting a sum of double exponential curve to the time-course of NPQ relaxation after transition from high (15 minutes of 2000 PPFD) to low light (50 minutes of 50 PPFD) (Dall’Osto et al. 2014; Long et al. 2022).

### Gas exchange measurements

During vegetative (V4-V5, 29-30 days after planting (DAS)) and reproductive (R5-R6, 93-94 DAS) stages, CO_2_ assimilation response curves to light were measured simultaneously with chlorophyll fluorescence following the protocol described by (De Souza et al. 2022) for NPQ activation and fluctuating light conditions. Theoretical maximum quantum efficiency of CO_2_ assimilation (ΦCO_2, max_) and theoretical maximum quantum efficiency of linear electron transport (ΦPSII_max_) were calculated as the slope of assimilation (A) over quanta and electron transport rate (ETR) over quanta from 50-200 µmol m^-2^ s^-1^. Non-photochemical quenching during low light (NPQ_low-light_), photochemical quenching (qP), Fv’/Fm’, and instantaneous quantum yield of linear electron transport (ΦPSII) were averaged over the same time points. Averages for non-photochemical quenching during high light (NPQ), ETR, and A were also calculated for the light intervals greater than 1800 PPFD during the fluctuating light protocol. Measurements were done on the youngest fully expanded leaves using a portable gas exchange system with a cuvette integrated with a modulated chlorophyll fluorometer and light source (LI-6800; Li-Cor, Lincoln, NE, USA).

The maximum apparent carboxylation rate by Rubisco (*V*_c,max_) and the maximum regeneration of ribulose-1,5-bisphosphate expressed as electron transport rate (*J*_max_) under steady-state were calculated from fitting A/Ci curves obtained using the LI-6800 as described above. For A/Ci curves, the light intensity was kept constant to 2000 µmol m^-2^ s^-1^ (90% red, 10% blue), block temperature at 28 °C, relative humidity inside the chamber at 60% ± 2%, and the reference CO_2_ concentration was varied following the sequence: 400, 300, 200, 150, 100, 75, 50, 400, 400, 600, 800, 1000, 1200, 1500, 1800 and 2000 µmol mol^-1^ for plants grown at a[CO_2_] after Long and Bernacchi, (2003) and 600, 400, 300, 200, 150, 100, 75, 50, 600, 600, 800, 1000, 1200, 1500, 1800, 2000 µmol mol^-1^ for plants grown at e[CO_2_] per (Ainsworth et al. 2007). Prior to fitting the curves, values were adjusted for temperature response to 25°C according to Bernacchi et al., (2001) and McMurtrie and Wang, (1993).

### Measurement of growth and development

Plant height and number of nodes were measured from 24 DAS until 85 DAS, when the plants reached the R5-R6 stage. Plant height was measured from the cotyledon scar to the growing point. The stages of soybean development were evaluated weekly starting at 8 DAS until the end of the experiment using the Fehr & Caviness, (1977) method. Leaf area index (LAI) was measured on 45, 50, 57, 63, 72, 78, 86, 93, and 107 DAS using a portable plant canopy analyzer (LAI-2200C; Li-Cor, Lincoln, NE, USA) at ground level between the middle two rows of each plot. Stem biomass and seed weight were determined after harvesting once the plants reached R8 stage of development. Harvest was done on October 7^th^, 2021 at 121 DAS. Ten plants of each plot were collected and placed in individual paper bags. Total number of pods, number of seed per pod, and stem biomass were determined after drying the plants in an oven at 60°C until constant weight. Harvest index was calculated as the ratio of seeds to the total above-ground biomass. Seed biomass per plot was determined by hand harvesting and threshing (LD350 Laboratory Thresher, Wintersteiger, Austria) all remaining plants per plot.

### RNA-SEQ analysis

Leaf samples were collected at 31 DAS and 86 DAS, during V4-V5 and R5-R6 stages, respectively, between 9:00 am and 11:00 am. Using a cork borer, three 13.4 mm leaf disks from the youngest fully expanded leaf of three different plants within a plot were sampled and pooled together. Samples were placed in 2mL nuclease-free tubes, immediately frozen in liquid nitrogen, and stored at -80°C. Samples were ground to a fine powder with 4 mm SPEX^TM^ stainless steel grinding beads (2150; SPEX^®^ SamplePrep, USA) using TissueLyser Universal Laboratory Mixer-Mill disruptor (85210; QIAGEN, Germany), at 24 Hz, two times of 1.5 min, stopping to cool cassettes in liquid N_2_.

Total RNA was extracted using the RNeasy Plant Mini Kit (74904; Qiagen, Germany) according to the manufacturer’s protocol with the following modification: RPE buffer wash was increased from one to 4-5 times. Further, the total RNA sample was digested with RNase-Free DNase Set (79254; Qiagen, Germany) to remove any traces of genomic DNA contamination. Total RNA concentration was initially evaluated using NanoDrop™ One Microvolume UV-Vis Spectrophotometer (Thermo Scientific™, USA). The quality of each RNA sample was determined in 1.2% agarose-TAE (0.5x) gel electrophoresis containing SYBR™ Safe DNA Gel Stain (Thermo Fisher, USA) by evaluating the integrity of the 28S and 18S ribosomal RNA bands and absence of smears. Construction of libraries and sequencing using the Illumina NovaSeq^TM^ 6000 system (Illumina, CA, USA) were performed at the Roy J. Carver Biotechnology Center at the University of Illinois at Urbana-Champaign. Purified total RNAs were run on a Fragment Analyzer (Agilent, CA, USA) to re-evaluate RNA integrity. The total RNAs were converted into individually barcoded polyadenylated mRNAseq (PolyA) libraries with the Kapa Hyper stranded mRNA Sample Prep kit (08098123702; Roche, IN, USA). Libraries were barcoded with Unique Dual Indexes (UDIs), which have been developed to prevent index switching. The adaptor-ligated double-stranded cDNAs were amplified by PCR for 8 cycles with the Kapa HiFi polymerase (07958927001; Roche, IN, USA). The final libraries were quantified with Qubit (ThermoFisher, MA, USA), and the average cDNA fragment sizes were determined on a Fragment Analyzer. The libraries were diluted to 10 nM and further quantified by qPCR on a CFX Connect Real-Time qPCR system (Biorad, CA, USA) for accurate pooling of barcoded libraries and maximization of the number of clusters in the flowcell.

The barcoded RNA-SEQ libraries were loaded on two SP lanes for 101 cycles on a NovaSeq^TM^ 6000 sequencing system. The libraries were sequenced from one end of the fragments for a total of 100bp. The Fastq read files were generated and demultiplexed with the bcl2fastq v2.20 Conversion Software (Illumina, CA, USA). Raw reads were deposited in the NCBI Sequence Read Archive, BioProject accession GSE270020. Reads from 32 RNA-SEQ libraries were first filtered to remove low length reads (i.e., reads <34). The reads were aligned to soybean genome Wm82.a4.v1 (downloaded from Phytozome (Sreedasyam et al. 2023) Phytozome genome ID: 508, version: *Gmax_Wm82_a4_v1* (Valliyodan et al. 2019)) using STAR aligner at default parameters (Dobin et al. 2013). The sequence of the three arabidopsis VPZ genes (violaxanthin de-epoxidase (*AtVDE, AT1G08550*), photosystem II subunit S (*AtPsbS, AT1G44575*), and zeaxanthin epoxidase (*AtZEP, AT5G67030*) (genome version TAIR10, annotation version Araport11) were added to soybean genome before aligning reads to the soybean genome. Further, *featureCounts* program from subreads package was used to count the number of reads assigned to each gene (Liao et al. 2014). Reads with multi-align flag were assigned fractionally to each of the mapped loci.

Four replicates for each treatment-genotype combination were sampled from the four blocks/rings in the field. To remove batch effect associated with the blocks/rings, ComBat-seq program was used (Zhang et al. 2020). Differential gene expression (DGE) from V4-V5 stage and R5-R6 were analyzed separately. DGE analysis was conducted with limma package (Ritchie et al. 2015). The low count genes were filtered out and libraries were normalized for size using Trimmed Mean of M -values (TMM) normalization method (Robinson and Oshlack 2010). Counts were transformed to log2-counts-per-million (logCPM) and were further transformed using ‘voom’ transformation. Differential expression analysis was carried out by fitting a linear model. Design matrix was setup to evaluate the statistical significance of main effects (CO_2_ and line) and interaction effect (CO_2_ x line). False discovery rate was controlled using Benjamini and Hochberg method. Adjusted p values were obtained from ‘p.adjust’ function from stats package in R. For main effects genes with p-adjusted < 0.05 and log_2_ fold change (FC) >1 were considered to be statistically significant unless stated otherwise. For interaction effect, genes with p-adjusted < 0.05 were considered significant. Genes significant for interaction term were clustered into co-expressed clusters to identify dominant interaction types. Clustering was performed using hierarchical clustering method. CO_2_ and line effect on predefined genes sets (KEGG pathways) was assessed using gene set enrichment analysis (GSEA). For both main effects the list of expressed genes was sorted based on the log_2_FC separately. WT and a[CO_2_] were considered the control condition for the analysis. ‘gseKEGG’ function of ClusterProfiler R package was used to run the analysis (Yu et al. 2012). Top 15 pathways with p-adjusted values <0.05 were considered significant.

A separate statistical analysis was conducted to test the effect of the growth stage, CO_2_ and line on the native and transgenic *ZEP*, *VDE* and *PsbS* genes. For this analysis, samples for two growth stages were analyzed together. DGE analysis was performed using the steps described above. Model design consisted of three main effects (the growth stage, CO_2_ and line) and their interactions. Raw p-values obtained for each main effect and interaction effect were corrected for multiple tests using Benjamini-Hochberg procedure as described above. Log_2_FC was calculated for each treatment, growth stage and line group. Genes with P adjusted value <0.05 and Log_2_FC > 1 were considered statistically and biologically significant.

### Statistical analysis

Data from physiological measurements were analyzed using the JMP® Software (JMP® Pro 15.0.0, SAS Institute Inc.) or R software (R Core Team 2025). Normal data distribution was checked using the Shapiro-Wilk W test, and equal variances using Brown-Forsythe and Levene tests (Dag et al.; Levene 1960; Shapiro and Wilk 1965; Brown and Forsythe 1974; Royston 1982). Significant differences were determined using a mixed model with restricted maximum likelihood (REML) considering [CO_2_] treatment, line, and the interaction between [CO_2_] and line as fixed effects, and block as a random effect (Cheung 2013). For time series measurements or measurements taken in different stages of development, the dates or stages were considered repeated measurements. Where the assumption of normality was not met, the non-parametric Kruskal-Wallis (K.W.) test was used (Hollander et al. 2015). Significant differences were considered when P≤0.1. That significant threshold was used to minimize the possibility of a type II error given the low replication level (*n*=4).

## RESULTS

### Faster NPQ relaxation and higher photosynthetic efficiency of VPZ-34A soybean is not affected by elevated [CO_2_] but is affected by growth stage

The time constants related to the fast (τ_1_; qE) and intermediate (τ_2_; qZ) relaxing components in VPZ-34A were significantly lower than the WT in both [CO_2_] conditions (Figure 1). While e[CO_2_] did not affect τ_1_ and τ_2_ in VPZ-34A, it reduced τ_1_ in WT by 49% and increased τ_2_ by 31% (Figure 1A). Maximum NPQ was higher in the VPZ-34A and remained unaffected by [CO_2_] (Figure 1C).

**Figure 1.**
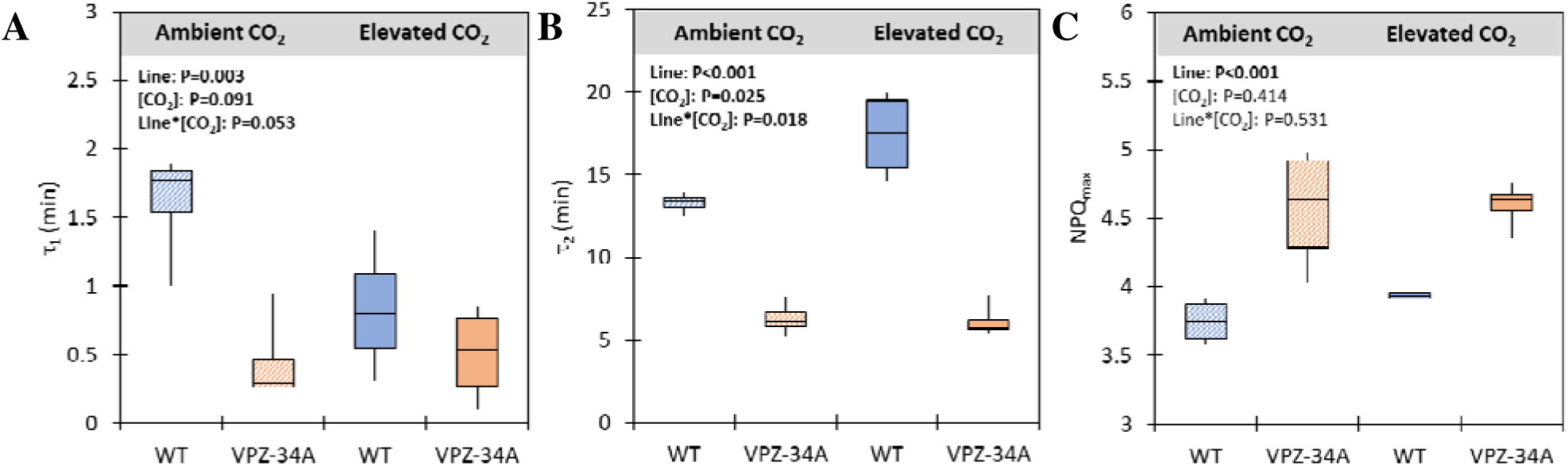
Time constants for **(A)** fast (τ_1_) and **(B)** medium (τ_2_) NPQ components, and **(C)** maximum NPQ (NPQ_max_) of WT and VPZ-34A grown at SoyFACE, Champaign, IL, USA under ambient and e[CO_2_]. Significant differences between lines, [CO_2_] and interaction line*[CO_2_] are in bold and indicated by P<0.1 (*n*=4).

Under fluctuating light, φPSII_max_ and NPQ_low-light_ were statistically different between VPZ-34A and WT only in the vegetative stage. φPSII_max_ in VPZ-34A was, on average, 7% and 5% greater than in WT at stages V4-5 and R5-6, respectively (Figure 2C & D). NPQ_low-light_ in VPZ-34A was 15% lower than WT at V4-5 and it dropped to 10% lower than WT at R5-R6 (Figure 2E & F). No significant differences in φCO_2, max_ were observed between genotypes (Figure 2A & B). Both genotypes exhibited reduced NPQ_low-light_ and increased φPSII_max_ under e[CO_2_] compared to those grown at a[CO_2_]. At e[CO_2_], φPSII_max_ for both WT and VPZ-34A was higher than at a[CO_2_] with a 4% gain at V4-5 and a 7% gain at R5-6 (Figure 2C & D). NPQ_low-light_ also showed greater improvement under e[CO_2_] with an 8% reduction in V4-5 and 11% reduction at R5-6 as compared to a[CO_2_] (Figure 2E & F). However, there was no significant interaction between VPZ-34A and e[CO_2_] for any of those measurements. Under NPQ activation, no significant changes in φPSII_max_ or φCO_2, max_ between VPZ-34A and WT were observed (Supplementary Fig. S3A-D). However, a significant difference in φPSII_max_ between ambient and elevated [CO_2_] were observed with a 3% reduction in e[CO_2_] at V4-5 and a 3% increase in e[CO_2_] at R5-6 (Supplementary Fig. S3C-D). φCO_2, max_ also showed a treatment effect with a 7% loss at e[CO_2_] at V4-5 and a line by treatment effect at R5-6 with a 10% increase in φCO_2, max_ for VPZ-34A at e[CO_2_] and a 10% decrease for WT at e[CO_2_] (Supplementary Fig. S3A-B). NPQ_low-light_ values were 13% lower in VPZ-34A in both [CO_2_] across both growth stages (Supplementary Fig. S3E-F). There was no significant line effect of line on mean assimilation during high light in fluctuating light at either growth stage (Supplementary Fig. S7E). There was no significant change in *V*_c,max_ and *J*_max_ between WT and VPZ-34A, but seasonal average of *V*_c,max_ was 23% lower in WT and 14% lower in VPZ-34A at e[CO_2_] as compared to a[CO_2_] (Table 1).

**Figure 2.**
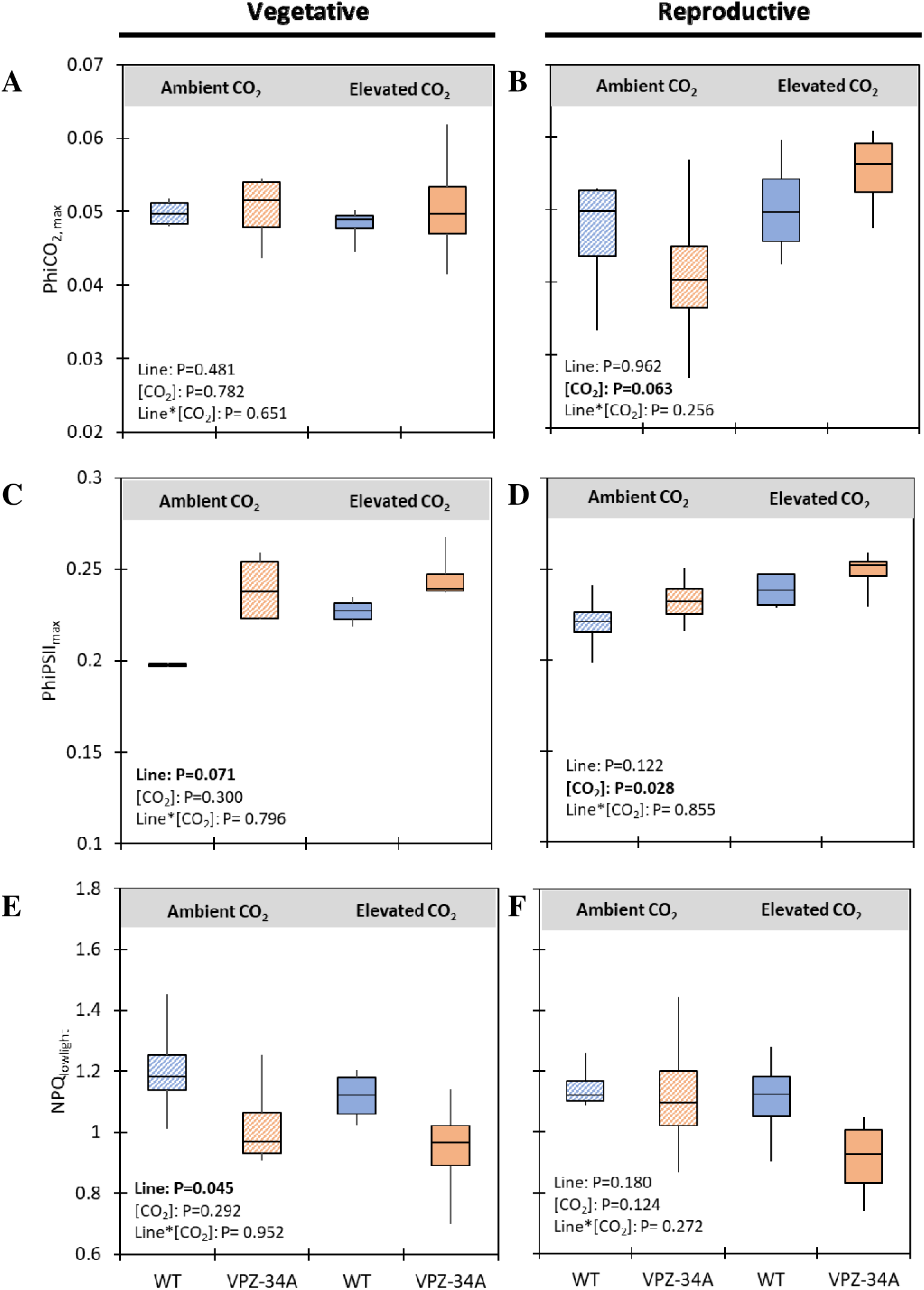
Maximum quantum efficiency of CO_2_ assimilation (ΦCO_2, max_) **(A-B)**, maximum quantum efficiency of linear electron transport (ΦPSII_max_) **(C-D)**, and non-photochemical quenching during sun to shade transitions (NPQ_lowlight_) **(E-F)** under fluctuating light of WT and VPZ-34A grown at SoyFACE, Champaign, IL, USA under ambient and elevated **a**[CO_2_]. Values represent repeated measurements from vegetative (V4-V5) **(A, C, E)** and reproductive (R5-R6) **(B, D, F)** stages. Significant differences between lines, [CO_2_] and interaction line*[CO_2_] are in bold and indicated by P<0.1 (*n*=4).

**Table 1.** Maximum rate of carboxylation by Rubisco (*V_cmax_*; µmol m^-2^s^-1^) and maximum rate of photosynthetic electron transport (*J_max_*; µmol m^-2^s^-1^) at steady-state in WT and VPZ-34A grown at SoyFACE, Champaign, IL, USA under ambient and e[CO_2_]. Values represent repeated measurements from vegetative (V4-V5) and reproductive (R5-R6) stages. Values are mean ± SE (*n*=4). Significant differences between lines, [CO_2_] and interaction line*[CO_2_] are in bold and indicated by P<0.1.

|  | Ambient CO<sub>2</sub> |  | Elevated CO<sub>2</sub> |  | ANOVA results |  |  |
| --- | --- | --- | --- | --- | --- | --- | --- |
| Season Averages |  |  |  |  |  |  |  |
|  | WT | VPZ-34A | WT | VPZ-34A | Line | [CO<sub>2</sub>] | Line*[CO<sub>2</sub>] |
| $V_{cmax}$ | 70.47 ± 6.57 | 66.08 ± 5.91 | 54.24 ± 5.96 | 57.79 ± 4.95 | 0.970 | **0.031** | 0.465 |
| $J_{max}$ | 125.99 ± 7.52 | 122.59 ± 9.67 | 115.21 ± 11.19 | 126.24 ± 7.83 | 0.691 | 0.703 | 0.443 |
| Vegetative (V4-V5) |  |  |  |  |  |  |  |
| $V_{cmax}$ | 83.08 ± 5.37 | 70.95 ± 5.23 | 59.18 ± 7.17 | 62.32 ± 6.10 | 0.501 | **0.033** | 0.269 |
| $J_{max}$ | 126.48 ± 5.38 | 117.61 ± 7.38 | 100.68 ± 9.71 | 122.34 ± 9.69 | 0.458 | 0.233 | **0.097** |
| Reproductive (R5-R6) |  |  |  |  |  |  |  |
| $V_{cmax}$ | 57.85 ± 8.15 | 61.21 ± 10.95 | 49.30 ± 9.89 | 53.25 ± 7.98 | 0.687 | 0.378 | 0.974 |
| $J_{max}$ | 125.50 ± 15.31 | 127.58 ± 19.12 | 129.74 ± 18.7 | 130.13 ± 13.5 | 0.945 | 0.851 | 0.967 |

### The intensity of plant growth stimuli and development response to elevated [CO_2_] is similar between WT and VPZ-34A

At 85 DAS, plants grown under e[CO_2_] were 14.6% taller than those under a[CO_2_], with no significant difference between WT and VPZ-34A (Figure 3A). At e[CO_2_], WT plants had on average three more nodes than at a[CO_2_], while the VPZ-34A had on average only two more nodes (Figure 3B). While the growth stages did not differ between VPZ-34A and WT under ambient or e[CO_2_] during the vegetative stages (VE to V8), most VPZ-34A plants reached R8 stage at e[CO_2_] faster than WT (DAS 126; Figure 4).

**Figure 3.**
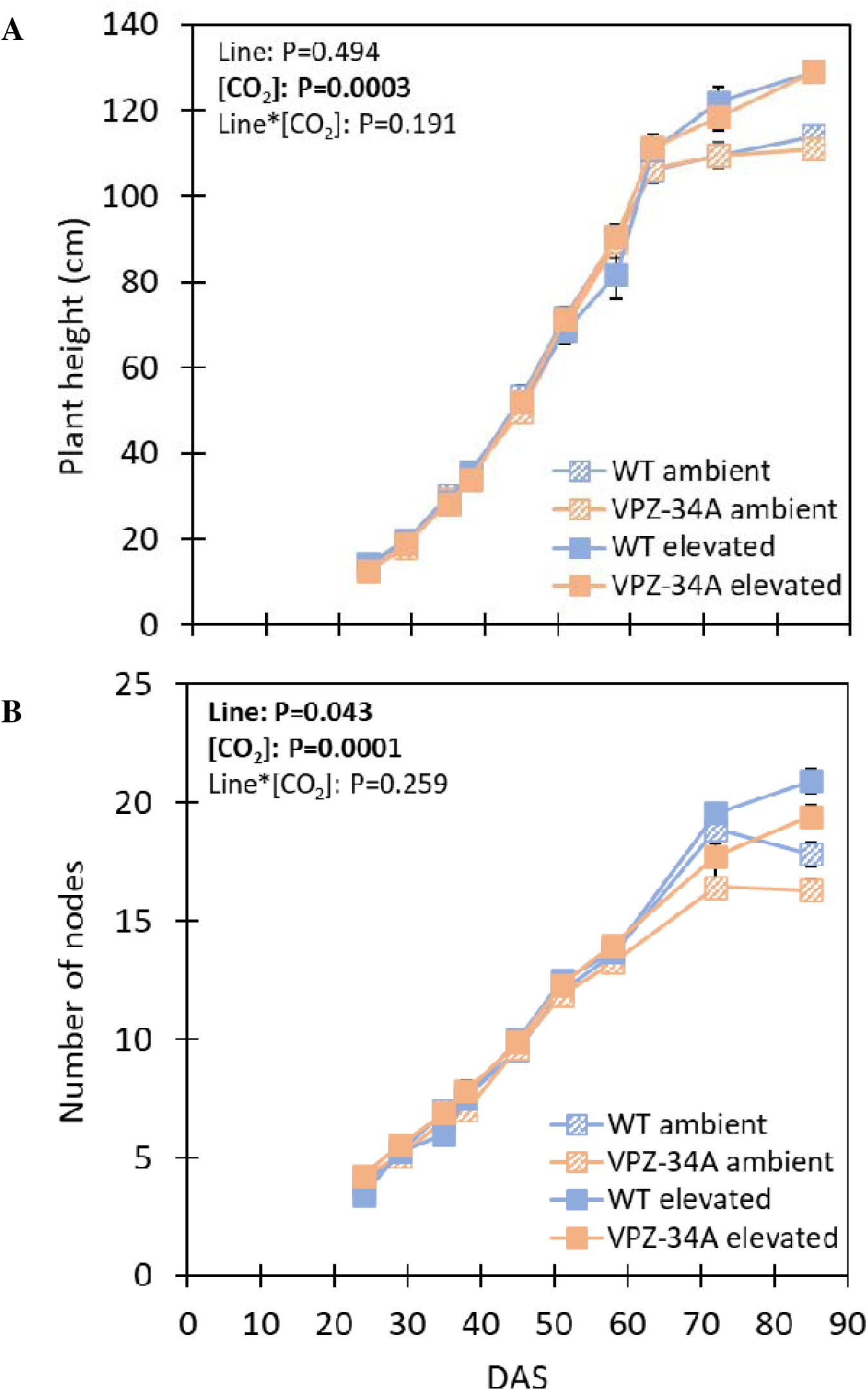
Plant height (cm) **(A)** and number of nodes **(B)** of WT and VPZ-34A grown at SoyFACE, Champaign, IL, USA under ambient and e[CO_2_]. Values are mean ± SE (*n*=4). DAS = days after sowing. Significant differences between lines, [CO_2_] and interaction line*[CO_2_] are in bold and indicated by P<0.1.

**Figure 4.**
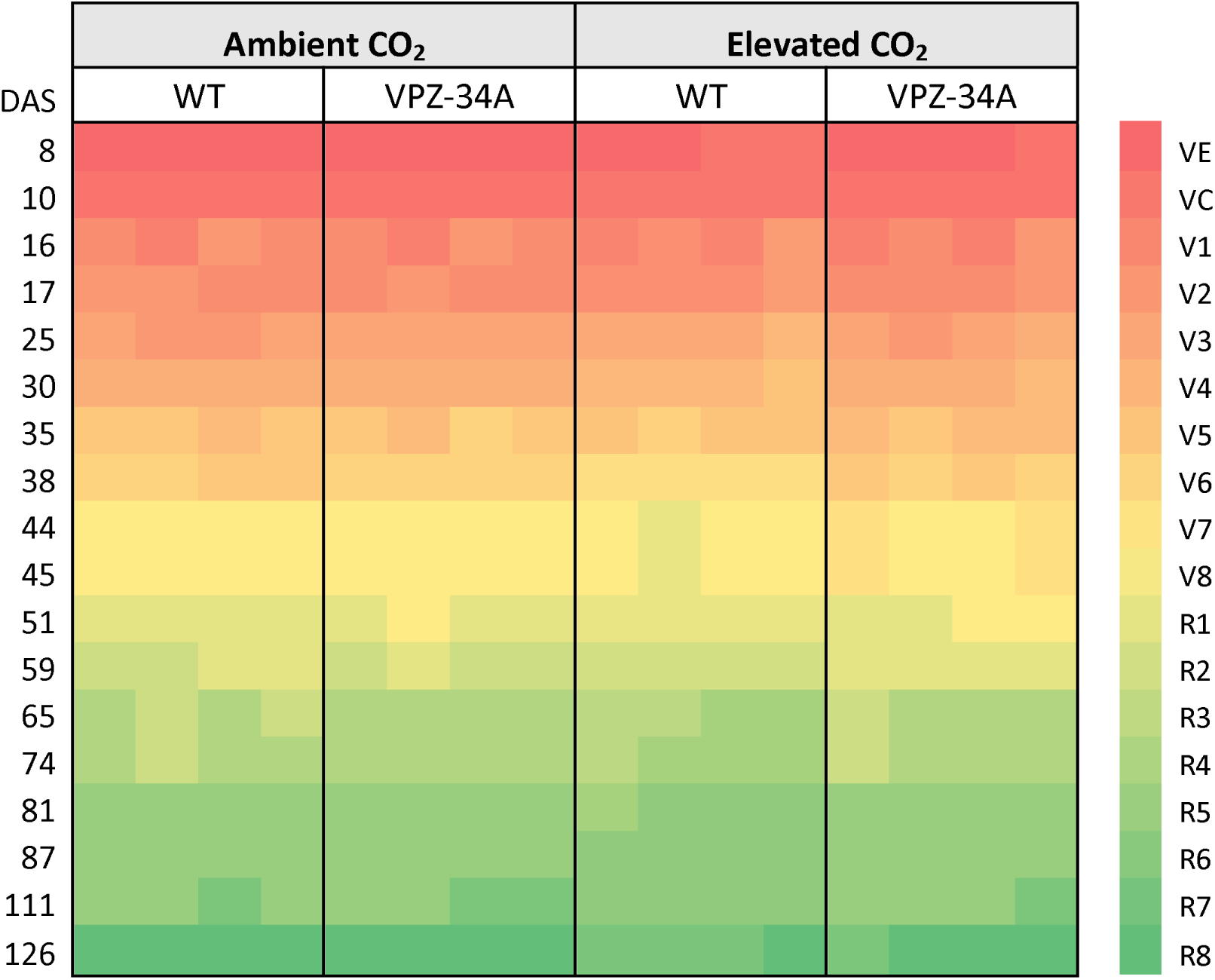
Developmental growth stages accessed from 8 days after sowing (DAS) to 126 DAS of wild-type (WT) and VPZ-34A grown at SoyFACE, Champaign, IL, USA under ambient and e[CO_2_]. Each column represents the data for one plot (*n*=4). VE = vegetative stage emergence; VC = vegetative stage cotyledon; V1-V8 = vegetative stage from the first fully developed trifoliate (V1) to eight nodes (V8); R1-R8 = reproductive stages.

At harvest, stem and seed biomass per plant were 50-55% greater at e[CO_2_] compared to a[CO_2_], with no significant differences between lines (Figure 5A-B). Similarly, seed production per plot was not altered between WT and VPZ-34A, but plots under e[CO_2_] produced significantly more seeds than in ambient a[CO_2_] (Figure 5C). Despite no differences in seed biomass per plant or per plot, the harvest index of VPZ-34A plants was 3% greater than that of WT (Figure 5D). Harvest index was not affected by [CO_2_] (Figure 5D). Seeds in the VPZ-34A were overall 8% smaller than the WT (Figure 5F).

**Figure 5.**
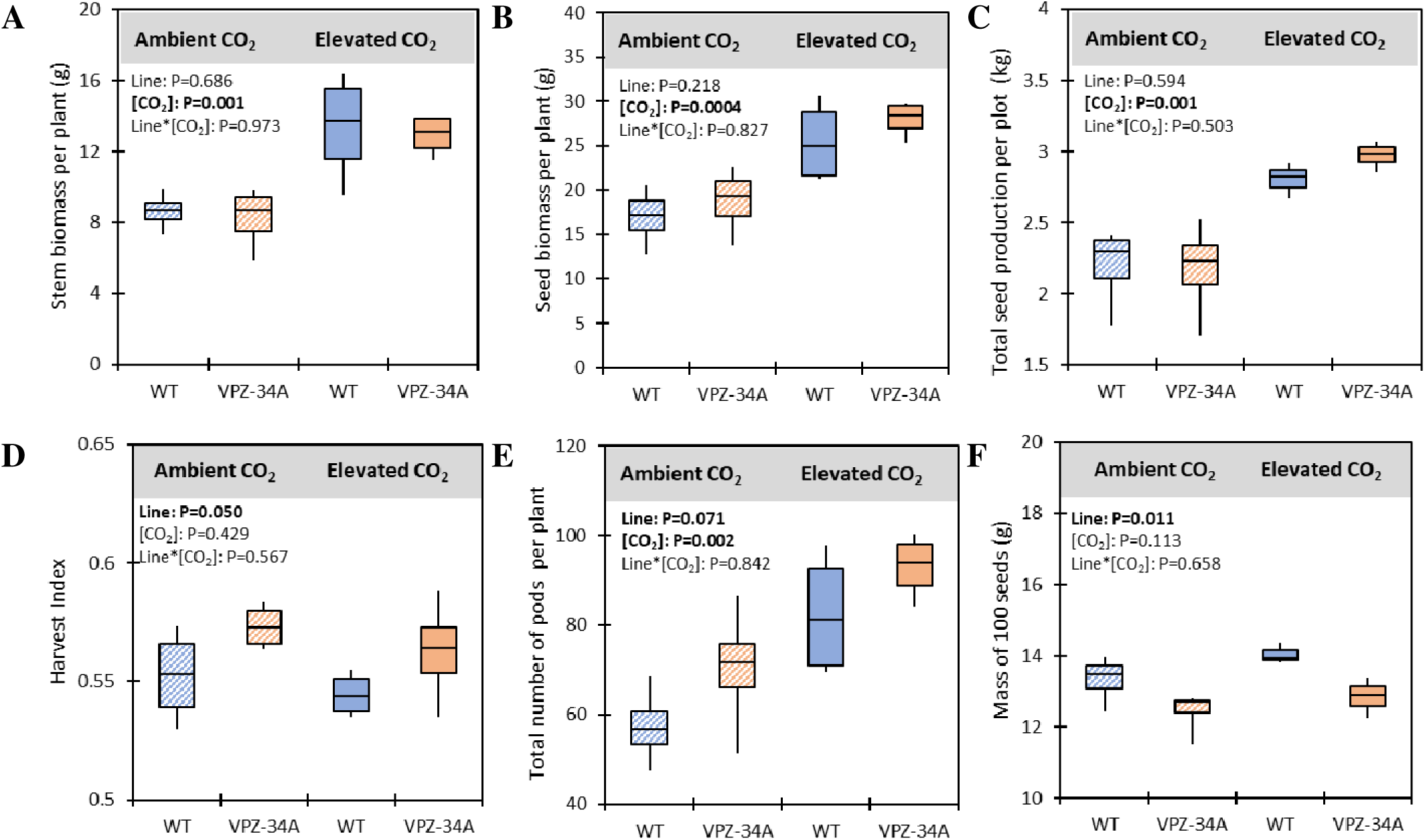
Stem **(A)** and seed **(B)** biomass per plant, seed biomass per plot **(C)**, harvest index **(D)**, total number of pods per plant **(E)**, and mass of 100 seeds **(F)** of WT and VPZ-34A grown at SoyFACE, Champaign, IL, USA under ambient and e[CO_2_]. Significant differences between lines, [CO_2_] and interaction line*[CO_2_] are in bold and indicated by P<0.1 (*n*=4).

### VDE, ZEP, and PsbS gene expression were significantly affected by the growth stage

*AtVDE*, *AtZEP* and *AtPsbS* were transcriptionally active in all transgenic samples. Their expression in the reproductive stage was generally lower than in vegetative stage. This growth stage effect was statistically significant for *AtVDE* and *AtZEP,* but not for *AtPsbS* (Supplementary Fig. S1, Supplementary File S1). However, growth under e[CO_2_] did not significantly affect the expression of transgenes. Gene expression of seven native VPZ genes (*GmZEP*, *GmVDE* and *GmPsbS*) was responsive to various independent variables and their interactions (Supplementary Fig. S1). However, only the growth stage resulted in a statistically significant and biologically meaningful expression changes (i.e. FDR < 0.05 and log_2_FC>1) for two *GmZEP* genes (*Glyma.09G00600* and *Glyma.11G055700*). Among the two *GmZEP* genes *Glyma.09G00600*’s expression was roughly eight times greater across both lines and [CO_2_] at the R5-R6 stage compared to V4-V5 stage. Upregulation of *Glyma.11G055700* at the R5-R6 stage ranged from one to four-fold.

### CO_2_ treatment and VPZ cassette on global transcription during vegetative and reproductive stages

For the main genotype effect at V4-V5 stage, 15 genes were identified as DEGs, while 35 genes were differentially expressed at the R5-R6 stage (FDR < 0.05 and log_2_FC>1) (Supplementary File S1). Notably, 13 of the 15 DEGs at from the V4–V5 stage were also differentially expressed at R6, indicating a consistent transcriptional signature associated with the VPZ cassette. Among the genes consistently differentially expressed across both stages, the most notable was the downregulation of *Glyma.08G087100* in VPZ-34A samples (Figure 6B & D). This gene is a homolog of a mitochondria localized thioredoxin *Trx o1* in arabidopsis. In addition, two ethylene responsive factors − *Glyma.09G248200* and *Glyma.18G244600* − which are homologous to the *BABY BOOM* (BBM) gene in arabidopsis were upregulated in VPZ-34A samples (Figure 6B & D). Given the small number of DEGs observed, gene set enrichment analysis (GSEA) was used to assess the entire list of expressed genes to delineate any weaker but pathway wide effects. GSEA revealed that flavonoid and isoflavonoid biosynthetic pathways were enriched at both growing stages for genotype effect (Figure 7A-B). In addition, enriched pathways at V4-5 included protein processing, N-glycan biosynthesis and photosynthesis antenna proteins (Figure 7A). Similarly, at R5-6, pathways such as carbon metabolism, oxidative phosphorylation, proteasome and glutathione metabolism were enriched. However, the range of log_2_FC among these pathways was less than one (Figure 7A-B). Thus, log_2_FC threshold for differentially expressed genes was relaxed to 0.5 to identify significant DEGs among the pathways identified by GSEA. At log_2_FC > 0.5, we identified 42 DEGs (38 upregulated and four downregulated in VPZ-34A) at vegetative stage and 168 DEGs (85 upregulated and 83 downregulated) at R5-6 stage. The relaxed log_2_FC cutoff allowed identification of three carbon metabolism related DEGs at R5-6, in addition to *Glyma.03G040600* (Pyrimidine 4, *PYD4*) which was significantly downregulated in VPZ-34A at log_2_FC>1 cutoff. *Glyma.01G091000* (phosphoenolpyruvate carboxylase 4, *PPC4*) was upregulated, *Glyma.16G204600* (Enolase 2, *ENO2*) and *Glyma.18G009700* (glyceraldehyde-3-phosphate dehydrogenase C subunit 1, *GAPC1*) were downregulated in VPZ-34A. Apart from carbon metabolism, glutathione metabolism at reproductive stage and N-glycan biosynthesis at vegetative stage were among the pathways identified via GSEA that consisted two and one DEGs respectively after log_2_FC cutoff relaxation.

**Figure 6.**
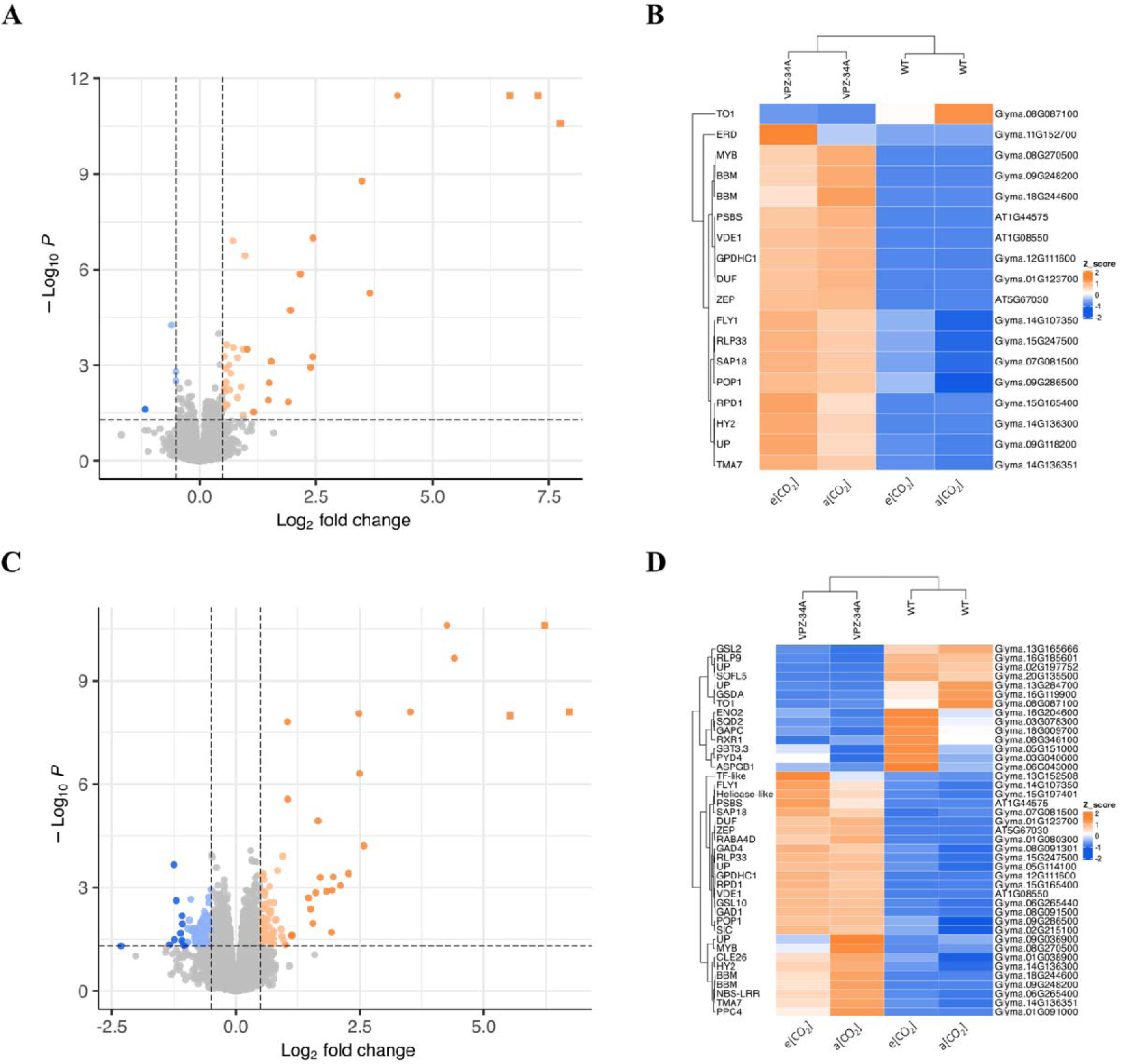
Main genotype effect differentially expressed genes (DEGs) at two growing stages. **(A & C)** Volcano plots representing DEGs between two lines at vegetative stage and reproductiv stage respectively. Circles represent soybean genes, squares represent transgenes. Darker shade of color represents significant genes with log_2_FC >1 **(B & D)** Heatmaps of significant DEG (log_2_FC>1) at vegetative stage and reproductive stage respectively

**Figure 7.**
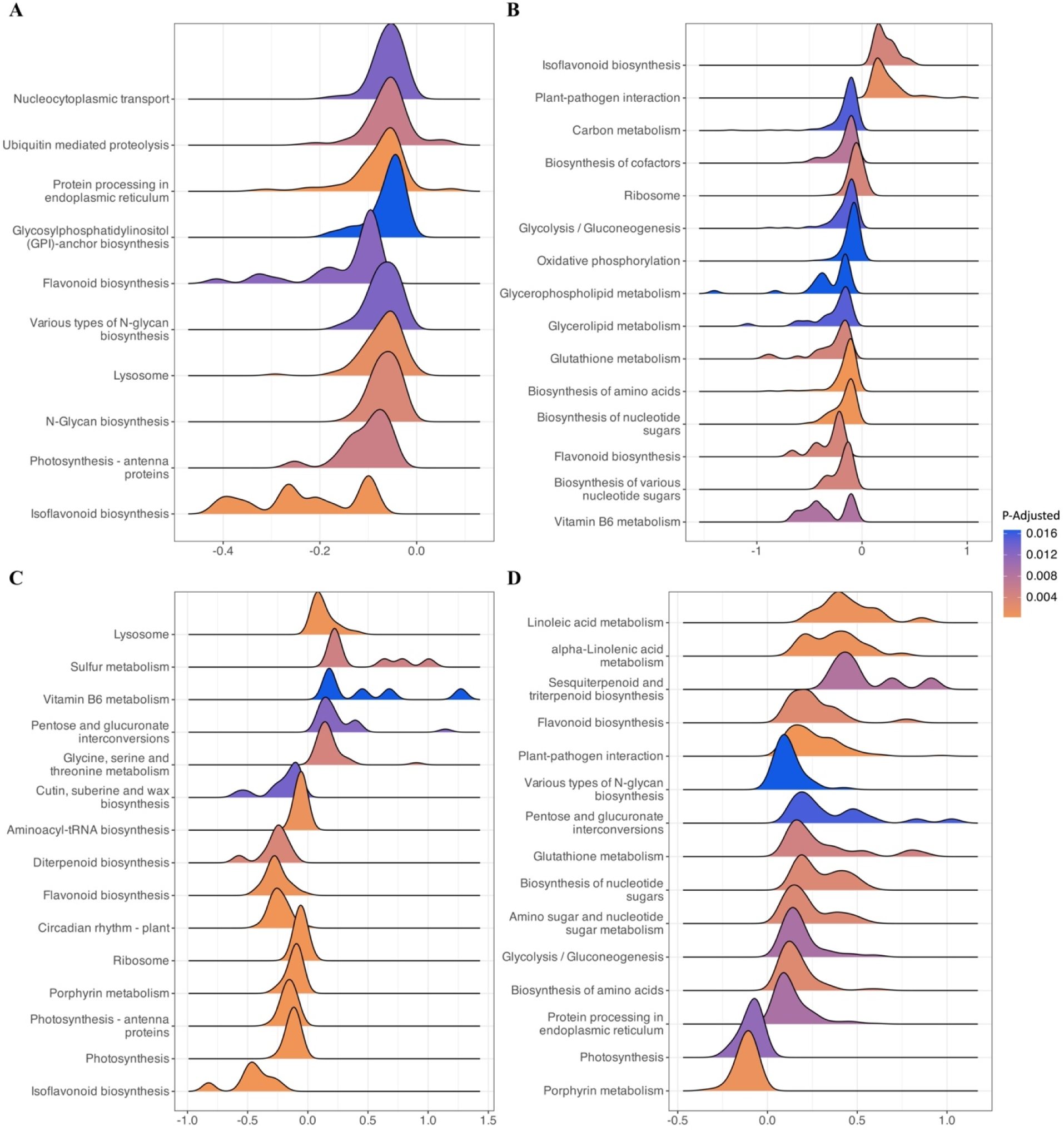
Gene set enrichment analysis (GSEA) of expressed genes at two growing stages using KEGG pathways as gene sets. **(A-B)** GSEA using log_2_FC calculated with VPZ-34A as treatment and WT as control at V4-V5 stage and R6 stage respectively. **(C-D)** GSEA using log_2_FC calculated with e[CO_2_] as treatment and a[CO_2_] as control at V4-V5 stage and R6 stage respectively. The color represents the statistical significance of the enriched pathway. The spread of the ridge represent the distributions of log_2_FC values of genes within that pathway.

Similar, to the genotype effect, a small CO_2_ effect was observed. At V4-V5 and at R5-6 stages respectively, 17 and 41 genes were differentially expressed in response to CO_2_ treatment (FDR < 0.05 and log_2_FC>1) (Supplementary File 1). The GSEA of all expressed genes revealed upregulation of sulfur metabolism, pentose and glucoronate interconversions, and glycine, serine and threonine metabolism at V4-V5 stage under e[CO_2_] (Figure 7C). The downregulated pathways under e[CO_2_] at V4-V5 stage included isoflavonoid biosynthesis, photosynthesis, ribosome and porphyrin metabolism (Figure 7C). At R6 stage, photosynthesis and porphyrin metabolism were found still downregulated, and linoleic acid, alpha-linolenic acid, plant-pathogen interaction, glutathione metabolism, amino acid metabolism and glycolysis were upregulated under e[CO_2_] (Figure 7D).

Lastly, genes responsive to interaction effect of line x CO_2_ were studied. At V4-V5 stage only 15 DEGs were significant for the interaction effect and clustered in four interaction clusters (Supplementary File S1, Supplementary Fig. S8). At R5-6 stage however, 283 genes were differentially expressed with top pathways being plant hormonal transduction, ribosome, spliceosome, RNA degradation, phagosome, oxidative phosphorylation and starch & sucrose metabolism as top pathways with genes greater than or equal to three. Genes significantly differentially expressed at R5-6 stage clustered into eight co-expression clusters (Supplementary File S1, Supplementary Fig. S9).

## Discussion

Using a previously characterized transgenic line (VPZ-34A), we set out to test the hypothesis that increases in light use efficiency, due to over-expression of *AtVPZ*, and carbon assimilation from e[CO_2_]_,_ would have an additive effect on seed production in soybean. However, we did not observe improvements in seed production or biomass in VPZ-34A relative to WT, in ambient or e[CO_2_]. Instead, stem and seed biomass, along with total seed production per plot, were only influenced by [CO_2_] (Figure 5 A-C). Importantly, despite seeing an increase in light use efficiency in VPZ-34A, we didn’t observe an increase in efficiency of carbon assimilation under fluctuating light, both in a[CO_2_] and e[CO_2_], which may underpin the lack of yield benefit.

The number of DEGs in response to *AtVPZ* was small (Figure 6), pointing to modest impact of *AtVPZ* on transcription of native soybean genes, although weak (low fold change) pathway wide changes were observed. The transcriptional response to e[CO_2_] in both lines is broadly consistent with previous reports for soybean (Ainsworth et al. 2006; Bredow et al. 2025). Although line specific CO_2_ effects were observed on gene expression, none of the growth phenotypes responded to the interaction effect. Therefore, the practical implications of such expression changes are unclear, therefore we have focused our discussion on the impact of *VPZ* on physiology and yield traits and related genes’ expression rather than the interaction with e[CO_2_].

### VPZ-34A transcriptional response to e[CO_2_] broadly similar to WT and previous FACE studies

In both VPZ-34A (and WT), the general transcriptional response to e[CO_2_] was consistent with previous studies. For example, upregulation of glycolysis at R5-6 is consistent with previous transcript analysis at SoyFACE (Leakey et al. 2009), as is a general downregulation of photosynthesis genes under e[CO_2_] Leakey et al. (2009) and Bredow et al. (2025). Similarly, changes in transcript abundance of genes belonging to stress response and lipid metabolism pathways were consistent with data described in Ainsworth et al. (2006) and Bredow et al. (2025). Therefore, the transcriptional response to e[CO_2_] is broadly preserved in both VPZ-34A and WT in the current study. Nonetheless, some of the responses to e[CO_2_] were different. For instance, we did not observe upregulation of protein degradation, or upregulation of respiratory pathways during vegetative stage as previously described by Ainsworth et al. (2006). These differences may be partly attributable to changes in environmental conditions between experiments, differences in techniques by which transcripts were evaluated (i.e. microarray vs. RNAseq), in the genotypes tested, and the time of day when samples were collected, as 21% of genes undergo circadian regulation (Locke et al. 2018).

### Potential reasons for lack of increase in seed production in VPZ-34A lines relative to WT

While the absence of an increase in seed production in VPZ-34A contrasts with one year of findings from De Souza et al. (2022), it is consistent with the second year (taken during the same year analyzed here). Sink limitation is commonly responsible for suppressing the benefits of increasing carbon assimilation (Ainsworth et al. 2002, 2011; Morgan et al. 2005; Bishop et al. 2015; Li et al. 2019), but it can be discounted as influencing the outcome, because we observed an increase in seed production in WT and VPZ-34A under e[CO_2_] (Figure 5C). Rather, harvest index and 100-seed weight were significantly impacted by VPZ expression, as previously seen for multiple VPZ events in soybean (De Souza et al. 2022). VPZ-34A produced a greater number of pods and seeds (Figure 5E & F), but a reduction in 100-seed weight offset any gains, resulting in a similar overall plot seed weight between lines under both ambient and e[CO_2_] (Figure 5C).

The observed effect of VPZ-34A on transcription of native soybean genes during reproduction was small. Grain filling happens during R5 stage, and by R6 seeds are already filed. Here samples for transcriptomics were taken at an intermediate stage (R5-R6) which may have impacted our ability to fully detect transcriptional changes impacting reproduction. However, there we did identify transcriptional changes that may have impacted yield. Particularly, stage specific downregulation of key carbon metabolism genes such as *GAPC1* and *ENO2* in VPZ-34A could have impacted the amount of photosynthate available for seed filling, which in turn, could be responsible for the smaller seeds in VPZ-34A plants (Figure 6C-D, 7B). Additionally, GSEA suggested pathway wide downregulation of carbon metabolism, but given the weak fold changes, it cannot be confidently concluded that the changes are biologically meaningful. How *AtVPZ* affects carbon metabolism in a growth stage dependent manner therefore remains to be understood.

Additionally, we found evidence for differential expression of two genes in VPZ lines with a potential role in seed development that could influence yield. *Glyma.09G248200* and *Glyma.18G244600* are homologs of arabidopsis *APETALA2 AP*2 domain containing protein PLT4/*BABY BOOM*. *AtBBM* regulates embryo and endosperm development, and knockout mutants of *AP2* gene are shown to have greater seed mass (Jofuku et al. 2005; Chen et al. 2022). The impact of lineage specific divergence of AP2 transcription factors is not fully understood (Jiang et al. 2020; Kerstens et al. 2020)), but upregulation of two putative developmental regulators as a result of *AtVPZ* expression combined with our observation of smaller seeds in VPZ-34A line warrants further investigation. An important caveat to this analysis is that we only analyzed transcriptional changes in leaves and not reproductive tissue, so it is unclear to what extent the observed transcriptional changes impact seed development. This may also be important as the transgenes themselves have the potential to impact seed development if expressed in reproductive tissues. For example, increased carotenoid content is correlated with decreased seed weight in chickpea and soybean (Abbo et al. 2005; Gebregziabher et al. 2022), and in arabidopsis, ZEP was identified as the major-effect locus for seed carotenoid content (Gonzalez-Jorge et al. 2016). In addition, expressing *AtVPZ* in reproductive tissues could affect NPQ and rates of pod photosynthesis, which have been shown to contribute to 14% in soy seed weight, and up to 25% in wheat and barley (Bort et al. 1994; Maydup et al. 2010; Cho et al. 2023). Analysis of gene expression atlases suggest that there is significant expression of *AtGAPA*, *AtFBA* and *AtRBCS* in embryonic, pod and seed tissues in arabidopsis, while a similar situation is true for homologous genes in soybean, indicating it is likely the transgenes are expressed to a significant level in reproductive tissues (Rhee et al. 2003; Winter et al. 2007; Almeida Silva et al. 2023).

Taken together, it cannot be ruled out that transcriptional changes induced by the VPZ cassette are impacting seed weight and pod number, through affecting carbon metabolism, developmental regulators or the transgenes themselves. It is therefore unclear whether impacts on seed weight can be decoupled from changes in leaf level photosynthesis, for example by restricting transgene expression to leaves. Nonetheless, our data provide several testable hypotheses for future research.

### Lack of observable increase in efficiency of CO_2_ assimilation in VPZ-34A

VPZ-34A was previously shown to possess increased theoretical quantum maximum light use efficiency (φPSII_max_) and increased theoretical quantum maximum efficiency of CO_2_ fixation (φCO_2, max_) under fluctuating light (De Souza et al. 2022). While we replicated the previously observed increase in φPSII_max_ in VPZ-34A, we did not see a corresponding increase in φCO_2, max_ (Figure 2A & C). Further, improvements in φPSII_max_ largely disappeared during reproductive stages (Figure 2D). This may potentially be attributable to the differential expression of the native *GmVPZ* genes, most notably an eight-fold increase in of *Glyma.09G000600* during reproductive stages, which is the ZEP homolog responsible for native LxL cycling (Leonelli et al. 2024) and may offset any gains from introducing the *AtZEP* in transgenic lines (Supplementary Fig. S1). In both tobacco and soybean, the lines and years with observed improvements in biomass correspond to improvements in φPSII_max_ and φCO_2, max_ (Kromdijk et al. 2016; De Souza et al. 2022). Without concurrent improvements in φPSII_max_ and φCO_2, max_, any gains in efficiency of light harvesting are not being used for CO_2_ fixation.

To explain why increasing φPSII_max_ may not lead to increased carbon assimilation, it is important to appreciate that φPSII_max_ is the product of two components: (1) the capacity for photochemical quenching (*F_q_’*/*F_v_*’; qP) and (2) light use efficiency (*F_v_’*/*F_m_’*) (Genty et al. 1989). Breaking down instantaneous φPSII at low light into these separate components reveals VPZ led to a decrease in NPQ in VPZ-34A, as seen by an increase in *F_v_’*/*F_m_’*, without impacting photochemical quenching (Supplementary Fig. S4E-G). Thus, while there were more electrons available during vegetative stages, they were not used for photosynthesis.

These extra electrons were likely used in alternate electron sinks such as cyclic electron flow (CEF), respiration, photorespiration, nitrogen assimilation, malate shuttle or Mehler reaction. Walker et al. (2014) previously implicated the malate-shuttle or Mehler reaction in balancing ATP/NADPH requirements under low light as opposed to CEF, and in accordance with these findings, we observed suppression of *Glyma.08G087100* in VPZ-34A, a putative mitochondrial thioredoxin o1 (*Trxo1*), which would be consistent with alterations in redox metabolism. In arabidopsis, Trxo1 is implicated in redox regulation of TCA cycle enzymes, alternative oxidase, and photorespiratory enzyme subunit glycine-decarboxylase (GDC) (Reinholdt et al. 2019), and knockout lines had perturbations to photorespiration and the malate shuttle pathways (Fu and Walker 2023; von Bismarck et al. 2023). Our measurement conditions were unable to provide insight into whether photorespiration is perturbed in VPZ-34A but provide an avenue for future research.

## Conclusion

Our data show that the overexpression of *VPZ* genes in soybean increases φPSII_max_ although it does not necessarily translate into increases in carbon assimilation and seed production. This is because there is a complex interplay between environment, xanthophyll cycle, photoprotection, rates of photosynthesis, stages of development and seed production, highlighted by changes in the overall gene expression in VPZ plants, and by the differential responses at the vegetative stages compared to reproductive stages. Several important questions remain: (1) how does *VPZ* expression impact seed weight, (2) what is the fate of reductant when φPSII is increased without impacting φCO_2_, (3) what is the impact of VPZ manipulation on *in situ* canopy photosynthesis, (4) if and how does VPZ expression perturb cellular redox regulation, and (5) precisely how do VPZ manipulations impact performance in response to abiotic stress. Unravelling the complexity identified in this study requires different approaches and further investigation and will be key for realizing the potential of VPZ expression as an engineering approach to increase crop yield.

## Data availability

Raw reads were deposited in the NCBI Sequence Read Archive (https://www.ncbi.nlm.nih.gov/sra), BioProject accession GSE270020.

## Acknowledgements

We dedicate this manuscript to the memory of Amy Marshall-Colón and Steve Long who started this work, for their vision, invaluable mentorship and genuine care for their co-workers. We are grateful for their oversight during the execution of the project from design, implementation, data collection, and preliminary data processing. It is with deep regret that we were unable to include them as authors as the final manuscript was prepared posthumously. Their shared enthusiasm and dedication to plant biology and improving food security through photosynthesis continues to inspire the authors to carry forward their vision.

In addition, we thank Anbarasu Karthikaichamy, Dhananjay Gotarkar, Meghan Burns, Jacob Milo and Emma Newlin for their assistance in sample collection, chlorophyll fluorescence imaging, and RNA extraction, Allison Altschuler for development scoring, node counts, and height, David Drag, Ron Eqdquilang, and Ben Thompson for field maintenance, and the SoyFACE research station staff and Dr. Lisa Ainsworth’s lab for installation and maintenance of the fumigation. This work was supported by the research project Realizing Increased Photosynthetic Efficiency (RIPE), funded from 2017-2023 under grant number OPP1172157 by the Gates Foundation, Foundation for Food and Agricultural Research, and the U.K. Government’s Department for International Development, and by Gates Agricultural Innovations grant investment ID 57248.

## Author Contributions

APS designed the research; APS, LD, SJB performed research; APS, LD and DS analyzed data; DS, LD, SJB and APS wrote the paper.

---

**Supplementary Figure 1.**
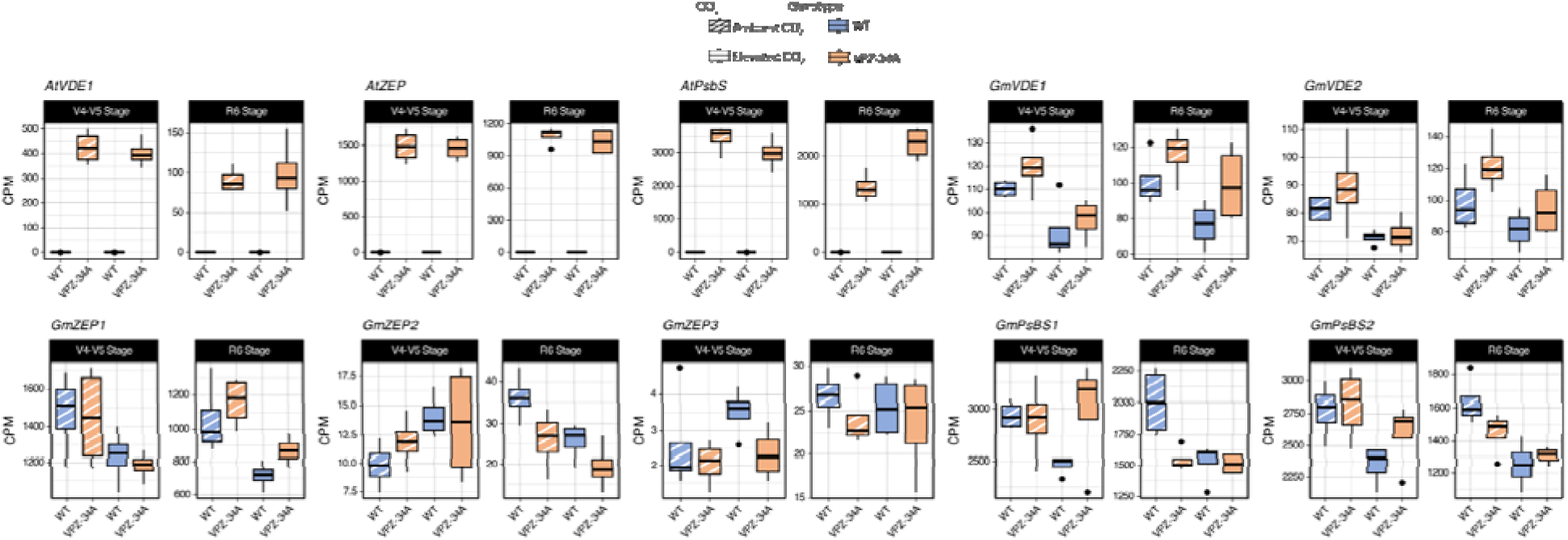
Gene expression of three *AtVPZ* transgenes and native VPZ gene across all treatment x stage combinations.

**Supplementary Figure 2.**
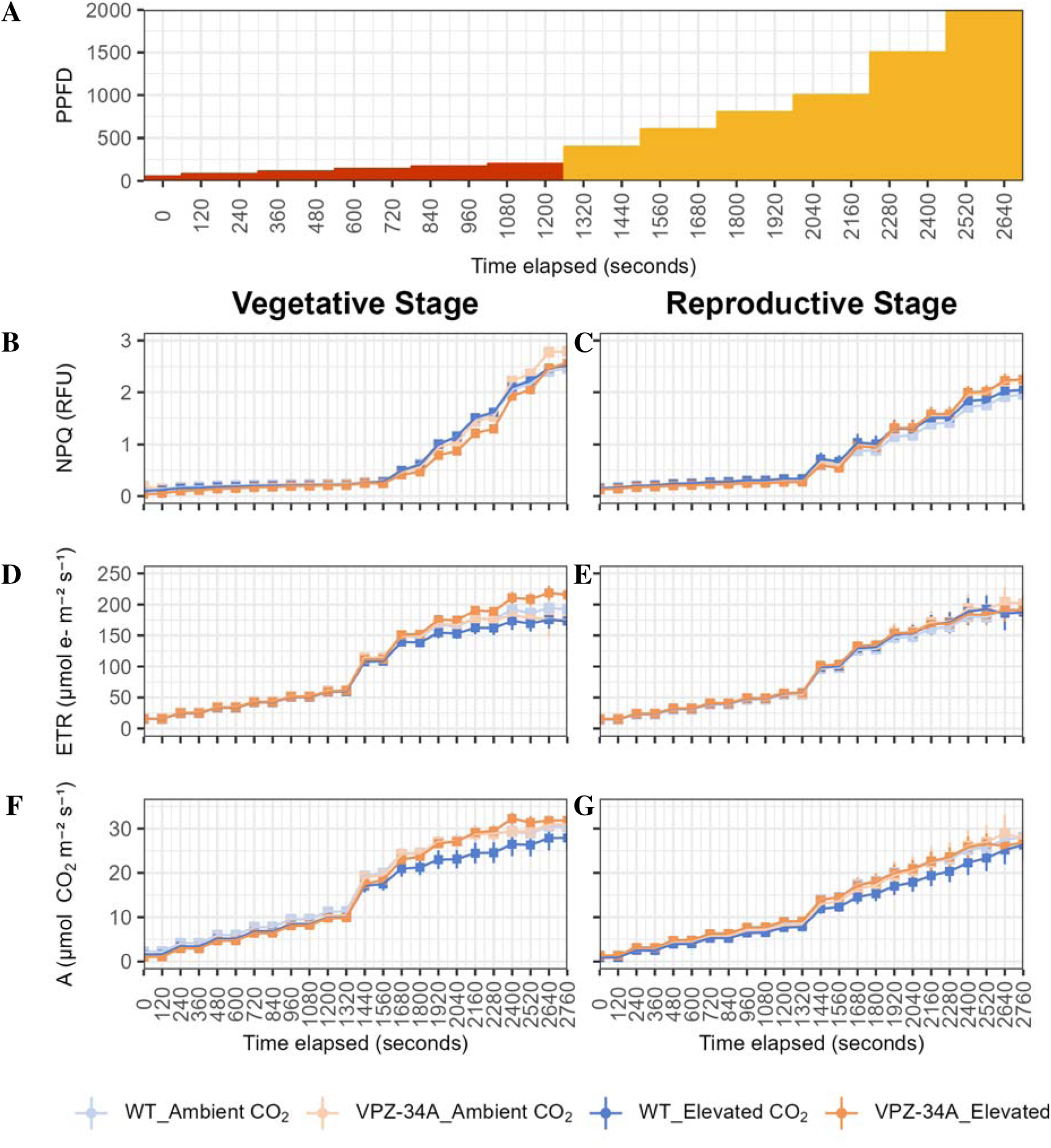
Light program for NPQ activation gas exchange protocol **(A).** Time points highlighted in red were used for calculations in SF3. Non-photochemical quenching (NPQ) **(B-C)**, electron transport rate (ETR) **(D-E)**, and carbon assimilation **(F-G)** over time elapsed during vegetative (V4-V5) (**B, D, F**) and reproductive (R5-R6) **(C, E, G)** stages of WT and VPZ-34A grown at SoyFACE, Champaign, IL, USA under ambient and [CO_2_].

**Supplementary Figure 3.**
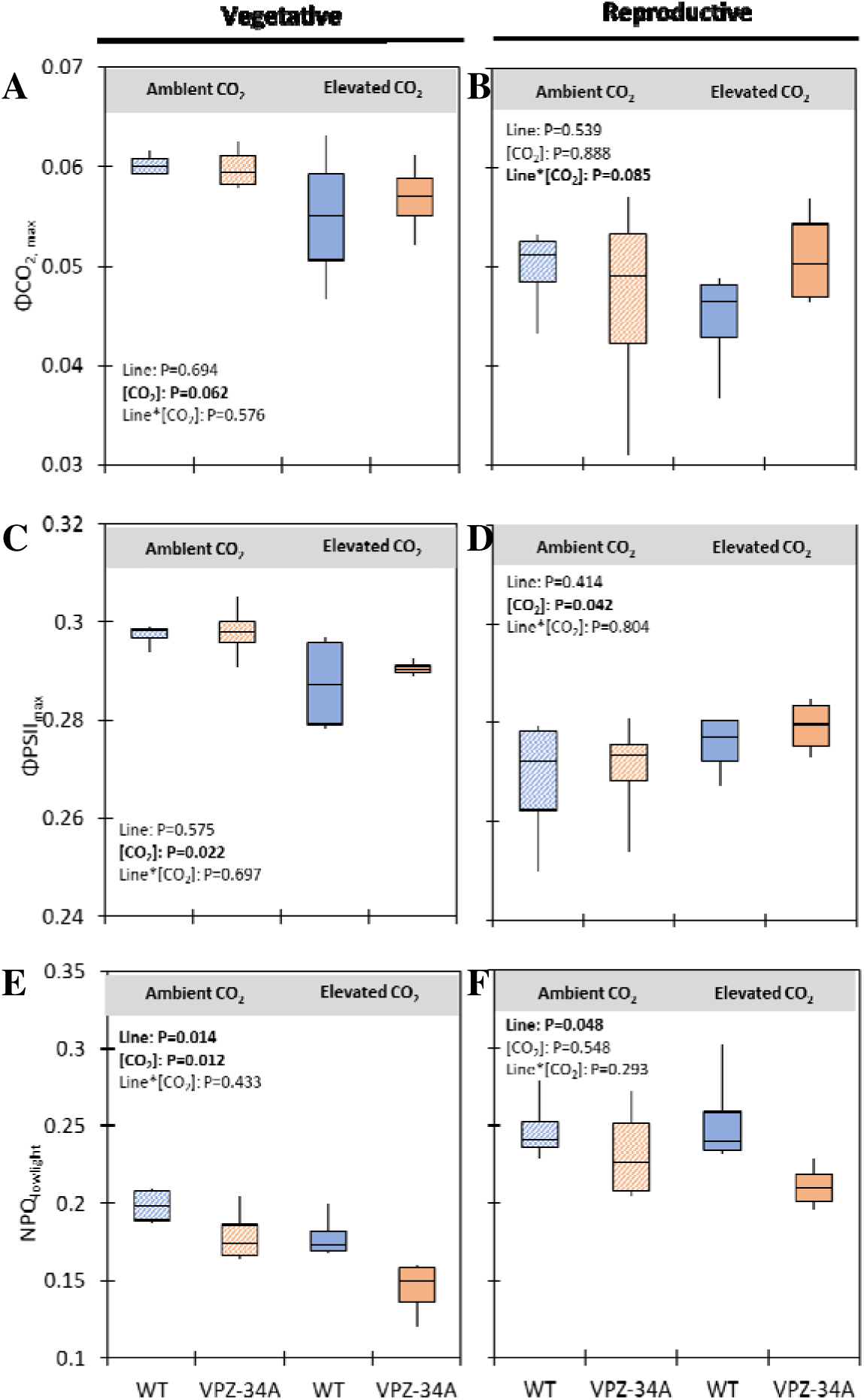
Maximum quantum efficiency of CO_2_ assimilation (ΦCO_2, max_) **(A-B)**, maximum quantum efficiency of linear electron transport (ΦPSII_max_) **(C-D)**, and non-photochemical quenching (NPQ_lowlight_) **(E-F)** during NPQ activation from 50-200 PPFD (SF2-A) of WT and VPZ-34A grown at SoyFACE, Champaign, IL, USA under ambient and e[CO_2_] for vegetative (V4-V5) **(A, C, E)** and reproductive (R5-R6) **(B, D, F)** stages. Values represent repeated measurements from vegetative (V4-V5) and reproductive (R5-R6) stages. Significant differences between lines, [CO_2_] and interaction line*[CO_2_] are in bold and indicated by P<0.1 (*n*=4).

**Supplementary Figure 4.**
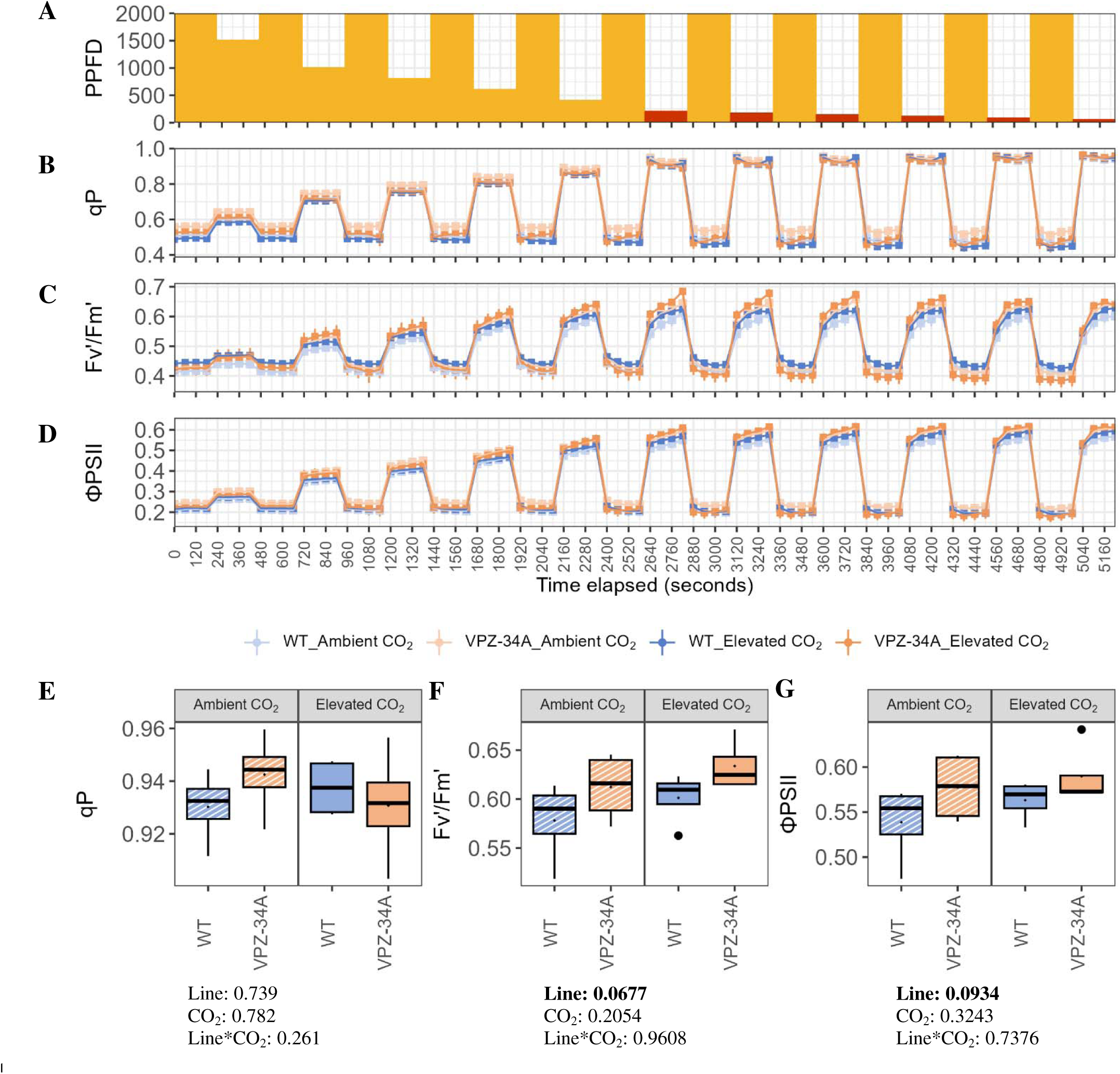
Light program for fluctuating light gas exchange protocol **(A).** Time points highlighted in red were used for calculations in E-G. Photochemical quenching (qP) **(B)**, Fv’/Fm’ **(C)**, and instantenous ΦPSII **(D)** over time elapsed during vegetative (V4-V5) stage. Means of qP **(E)**, Fv’/Fm’ **(F)**, and instantaneous ΦPSII **(G)** from low light (<220 PPFD) after high to low light transitions of WT and VPZ-34A grown at SoyFACE, Champaign, IL, USA under ambient and **e**[CO_2_]. Values represent repeated measurements from vegetative (V4-V5) stage. Significant differences between lines, [CO_2_] and interaction line*[CO_2_] are in bold and indicated by P<0.1 (*n*=4).

**Supplementary Figure 5.**
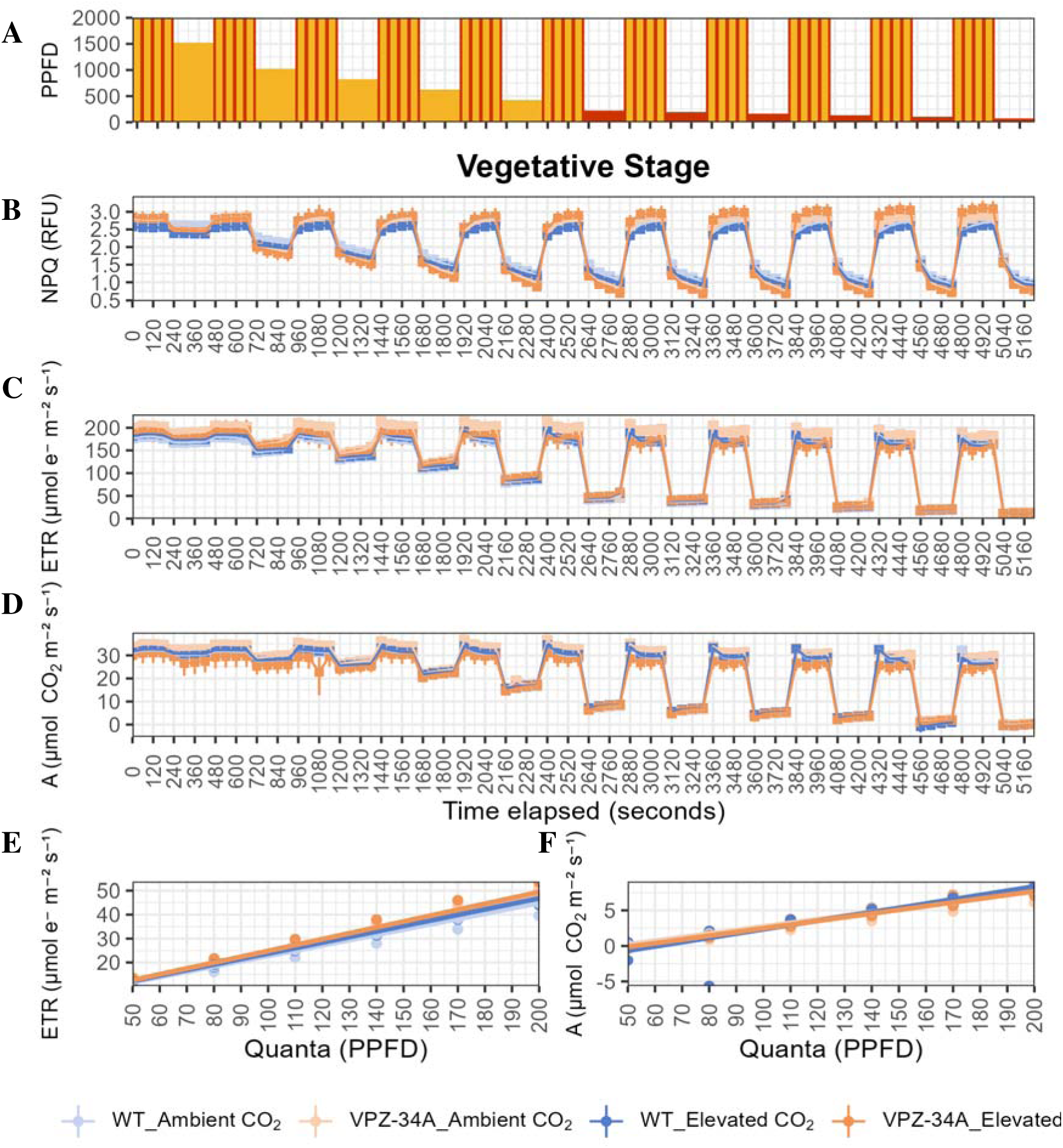
Light program for fluctuating light gas exchange protocol **(A).** Time points highlighted in red were used for calculations in E-F. Time points marked with red stripes were used for high light calculations (Fig. S6). Non-photochemical quenching (NPQ) **(B)**, electron transport rate (ETR) **(C)**, and carbon assimilation **(D)** over time elapsed during vegetative (V4-V5) stage. Mean linear model for slope calculations of theoretical maximum yield of photosystem II (ΦPSII_max_) **(E)** and theoretical maximum yield of carbon assimilation (ΦCO_2,max_) **(F)** calculated from 50-200 PPFD after high to low light transitions of WT and VPZ-34A grown at SoyFACE, Champaign, IL, USA under ambient and **e**[CO_2_]. Values represent repeated measurements from vegetative (V4-V5) stage. Significant differences between lines, [CO_2_] and interaction line*[CO_2_] for ΦPSII_max_ and ΦCO_2,max_ are shown in figure 2.

**Supplementary Figure 6.**
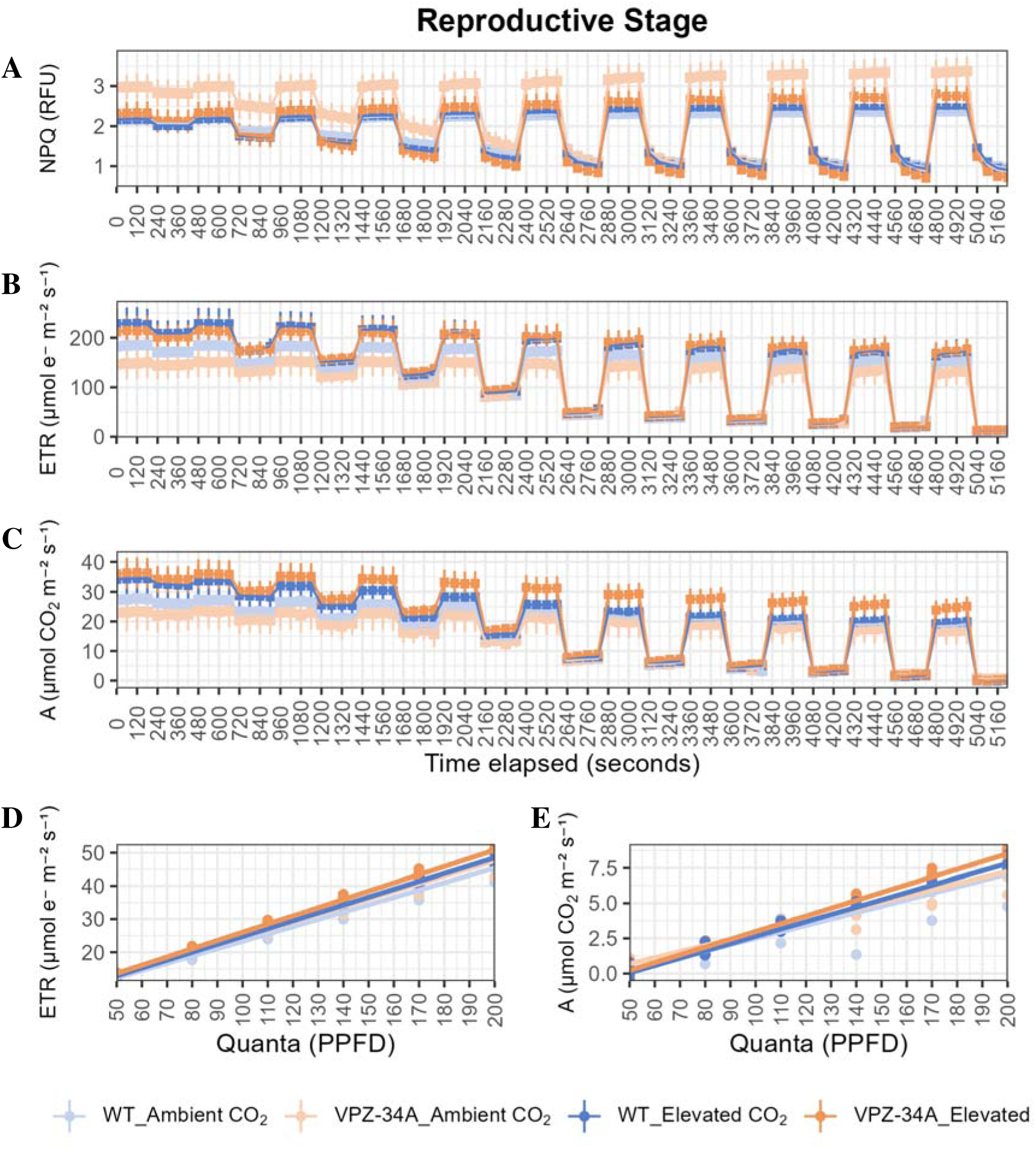
Non-photochemical quenching (NPQ) **(A)**, electron transport rate (ETR) **(B)**, and carbon assimilation **(C)** over time elapsed during reproductive (R5-R6) stage under the fluctuating light (Supplementary Fig. S5A). Mean linear model for slope calculations of theoretical maximum yield of photosystem II (ΦPSII_max_) **(E)** and theoretical maximum yield of carbon assimilation (ΦCO_2,max_) **(F)** calculated from 50-200 PPFD after high to low light transitions of WT and VPZ-34A grown at SoyFACE, Champaign, IL, USA under ambient and e[CO_2_]. Values represent repeated measurements from reproductive (R5-R6) stage. Significant differences between lines, [CO_2_] and interaction line*[CO_2_] for ΦPSII_max_ and ΦCO_2,max_ are shown in figure 2.

**Supplementary Figure 7.**
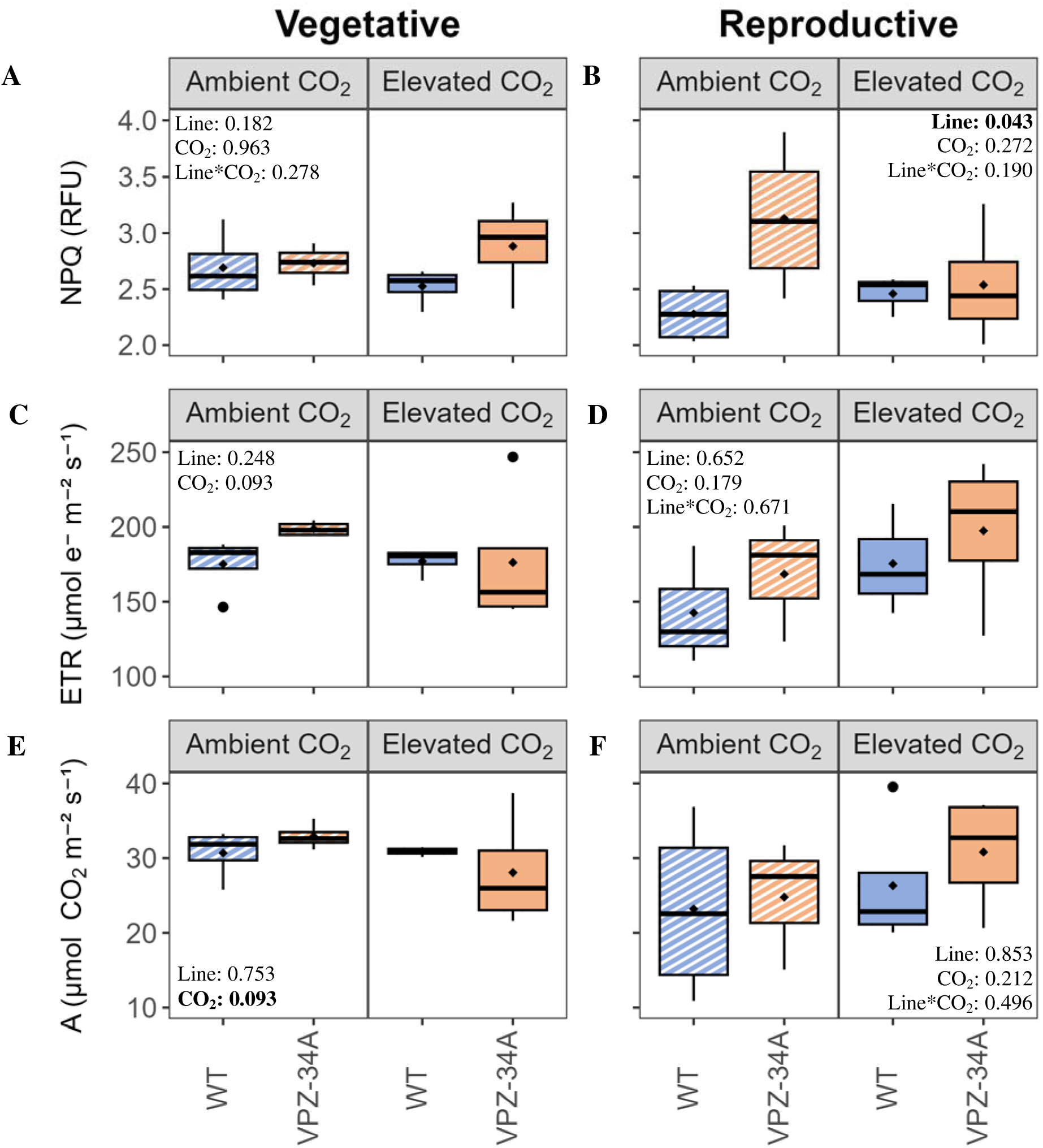
Mean non-photochemical quenching (NPQ) **(A-B)**, electron transport rate (ETR) **(C-D)**, and carbon assimilation (A) **(E-F)** at high light (>1800 PPFD) in fluctuating light (Supplementary Fig. S5A) during vegetative (**A, C, E)** and reproductive (**B, D, F)** stages in WT and VPZ-34A grown at SoyFACE, Champaign, IL, USA under ambient and **e**[CO_2_]. Significant differences between lines, [CO_2_] and interaction line*[CO_2_] for ΦPSII_max_ and ΦCO_2,max_ are in bold and indicated by P<0.1 (*n*=4). For vegetative mean ETR and A at high light, the assumption of normality was not met. The non-parametric Kruskal-Wallis (KW) test was used to evaluate line and [CO_2_] effects independently.

**Supplementary Figure 8.**
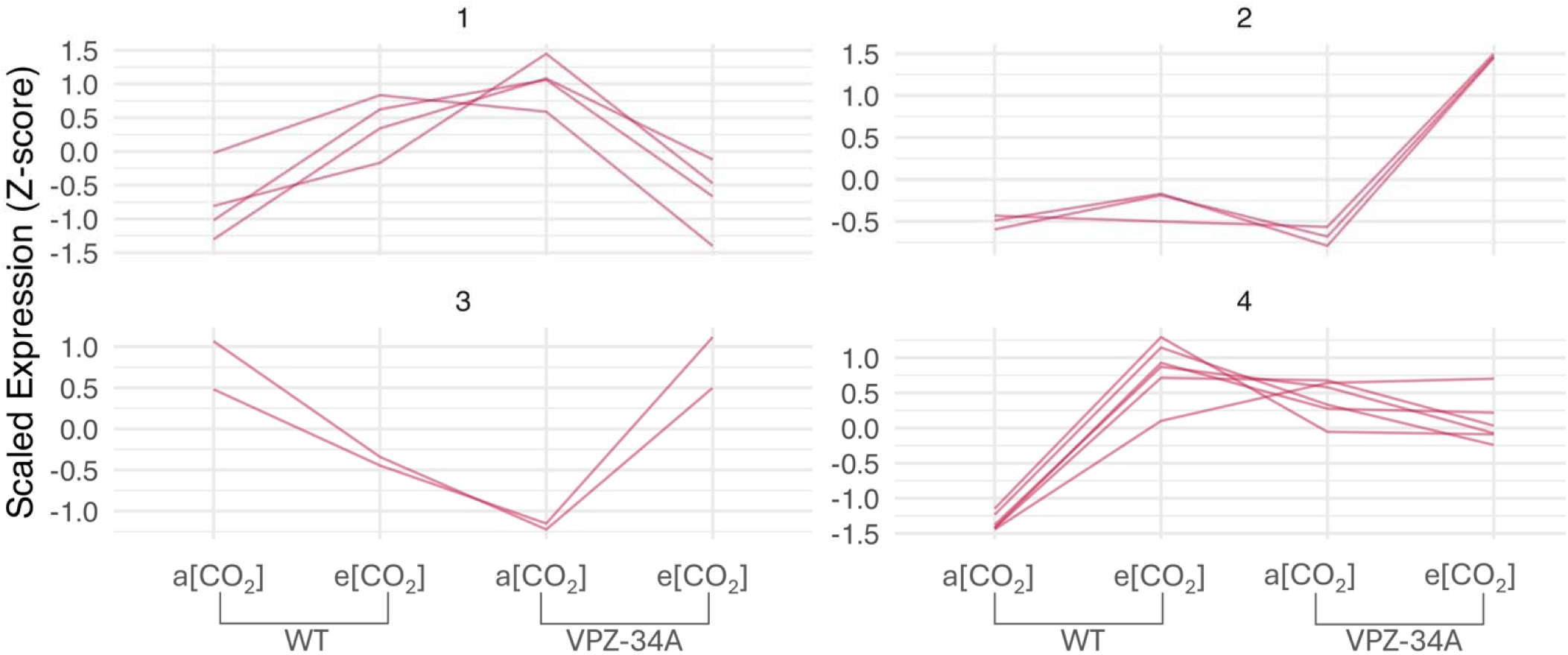
Co-expression clusters among the line x CO_2_ responsive DEGs at V4-V5 stage. Co-expression clusters were created using using hierarchical clustering of the differentially expressed genes.

**Supplementary Figure 9.**
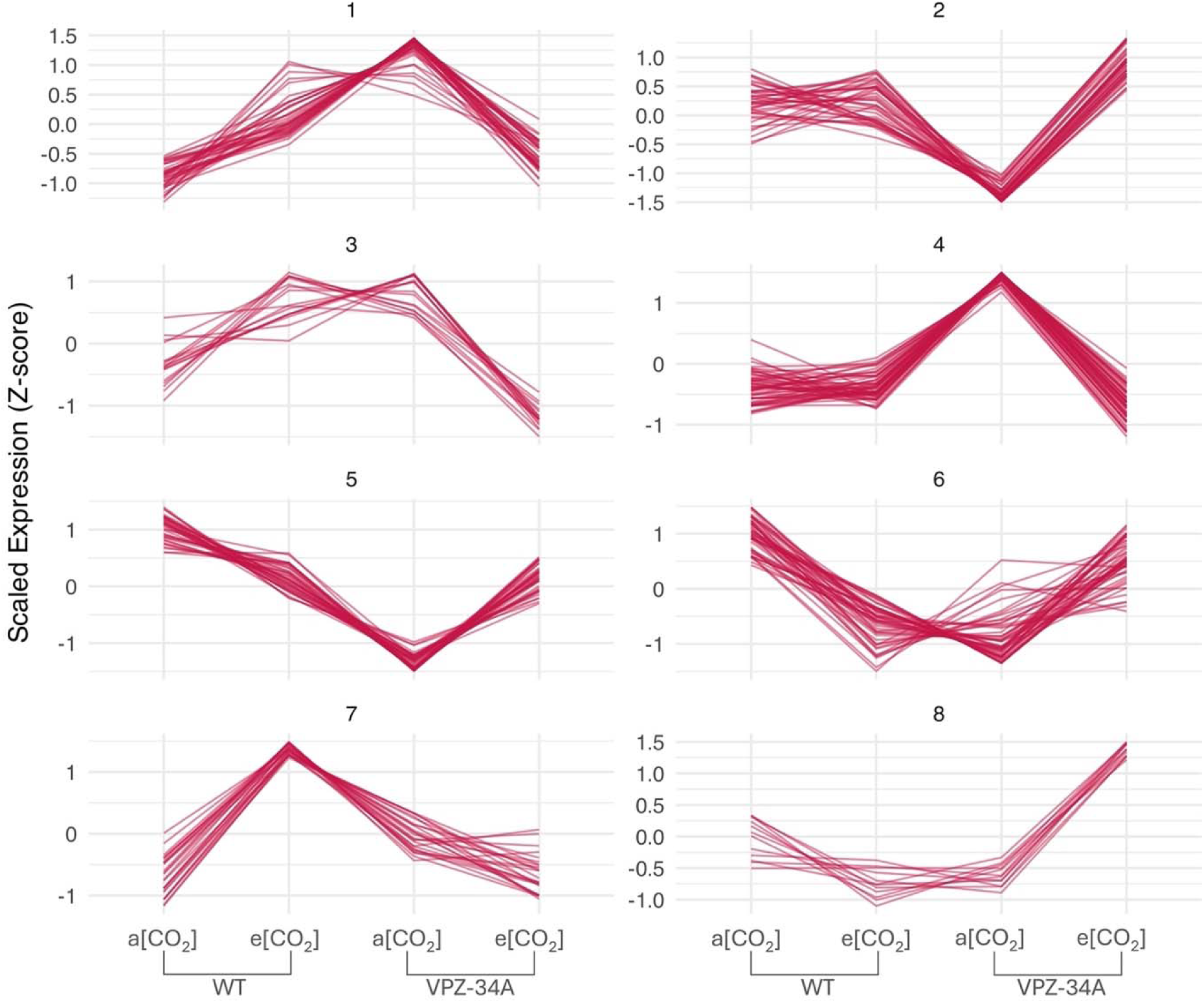
Co-expression clusters among the line x CO_2_ responsive DEGs at R6 stage. Co-expression clusters were created using using hierarchical clustering of the differentially expressed genes.

